# Selection constraints of plant adaptation can be relaxed by gene editing

**DOI:** 10.1101/2023.10.16.562583

**Authors:** Megan Ruffley, Ulrich Lutz, Laura Leventhal, Shannon Hateley, Wei Yuan, Jakob Keck, Seung Y. Rhee, Detlef Weigel, Moises Exposito-Alonso

**Author notes:** equal contribution (Megan Ruffley, Ulrich Lutz). Plant Resilience Institute, Departments of Biochemistry and Molecular Biology, Plant Biology, and Plant, Soil, and Microbial Sciences, Michigan State University, East Lansing, Michigan 48824, USA. Department of Integrative Biology, University of California Berkeley, Berkeley, CA 94720, USA. Howard Hughes Medical Institute, University of California Berkeley, Berkeley, CA 94720, USA. Institute of Physical and Theoretical Chemistry, University of Tübingen, 72074 Tübingen, Germany (Jakob Keck).

## Abstract

Climate change has already caused noticeable changes in species-wide traits, such as the well-documented acceleration of spring flowering. Because the evolutionary past has favored certain combinations of traits, some strategies like fast growth with early flowering that are adaptive today are at odds with other plant resilience strategies such as elevated water use efficiency. We know that the evolution of trait combinations is shaped by genomic constraints, but it is unclear whether and how this is affected by natural selection from climate change. Growing hundreds of *Arabidopsis thaliana* natural populations under different rainfall regimes revealed opposing natural selection on flowering time and water use efficiency, with strong antagonistic genetic correlations and contrasting causal alleles identified by Genome-Wide Association analyses. Inactivation of the central flowering regulator *FLC* in multiple, diverse accessions relaxed trait correlations in a genetic background-dependent manner and allowed for the emergence of a novel adaptive trait combination—early flowering and intermediate water use efficiency. Future climates are predicted to escalate conflicts in natural selection among adaptive traits, but our work shows that surprisingly simple genetic changes can help solve these conflicts.

---

Phenotypic trade-offs across individuals within a species, or across species, are common in nature ^1^. Trade-offs may arise as a result of local natural selection favoring certain trait combinations ^2,3^ or an overlap in the genetic loci involved in those traits, or both. These trade-offs may limit the ability of organisms to simultaneously optimize multiple traits in a new environment, as the most favorable trait combination may not yet exist, which will thus constrain adaptation ^4^. For example, plant phenology and ecophysiological traits relevant for adaptation in water limiting conditions—e.g., *escape* from drought periods through rapid reproductive cycling or *avoidance* of water loss through efficient CO_2_ uptake and maintenance of internal water pressure—often show strong trade-offs with each other ^5,6^. Given climate change predictions surpassing 2°C of global warming, with more frequent and extreme droughts ^7^, we are already observing an acceleration of phenological traits such as early spring flowering, which in turn is increasingly unsynchronized with pollinator emergence ^8–11^, as well as die-off of drought sensitive genotypes ^12^, suggesting evolutionary constraints preventing adaptation. We do not yet understand how correlated genomic trait architectures influence how natural selection acts on these phenotype combinations across environments, or whether such genetic trade-offs can be uncoupled. By combining trait data from *Arabidopsis thaliana* populations with field experiments, and using CRISPR/Cas9 to edit a major adaptive locus in multiple populations, we quantify a maladaptive trade-off between early flowering and water use efficiency (WUE), which may impede phenotype evolution due to a strong genetic constraint. While knocking down a central flowering regulator has the expected negative effect on WUE, it was very surprising that the correlation between flowering time and WUE is relaxed considerably, thus allowing natural selection to act directly to increase WUE.

To realize the phenotypic space of climatic adaptations in *A. thaliana*, we compared 64 ecologically-relevant traits related to life history and drought response in hundreds of geographically diverse accessions (see **Text SI, Fig. S1**). We determined the main axes of phenotypic variation with Principal Component Analysis (PCA) and overlaid a 2D color scheme in order to differentiate the suites of phenotypes by color (**Fig. 1A**) (**Text SII**). The main PC axis of variation captures the key phenotypic trade-offs of WUE and flowering time (highly correlated traits, n=246, Pearson’s *r* = 0.38, *p* < 5×10^-10^) (**Fig. 1B**), stomatal density and stomatal size ^13^, high and low seed dormancy ^14^, and a life history shift between rapid flowering without or only after vernalization ^15^; while the secondary axis is associated with germination rate (*i*.*e*. primary dormancy) and root architecture (**Fig. 1A**). The first axis describing within-species variation is significant and of general relevance, as it is also the first component structuring life history, lifespan, and physiological variation across hundreds of thousands of vascular plant species ^1^, often called a “fast versus slow” plant economic spectrum ^16–18^. PC1 also captures a well-characterized continuum in *A. thaliana* between investing resources in early reproduction (escapers) versus prioritizing WUE (avoiders) (**Fig. 1A**) ^19–21,22,23^. This gradient is apparent in trait correlations, with up to 40% of all traits involved in escape or avoidance (n=506; **Table S1**) being significantly correlated with flowering time (*i*.*e*., lifespan) ^24^, 17% with WUE ^25^, and 25% with growth rate ^18^ (**Table S3**). This indicates a massive overlap in traits related to adaptation, with substantial potential for shared genetic architectures that could constrain population adaptation phenotypic responses to new climates.

**Fig. 1.**
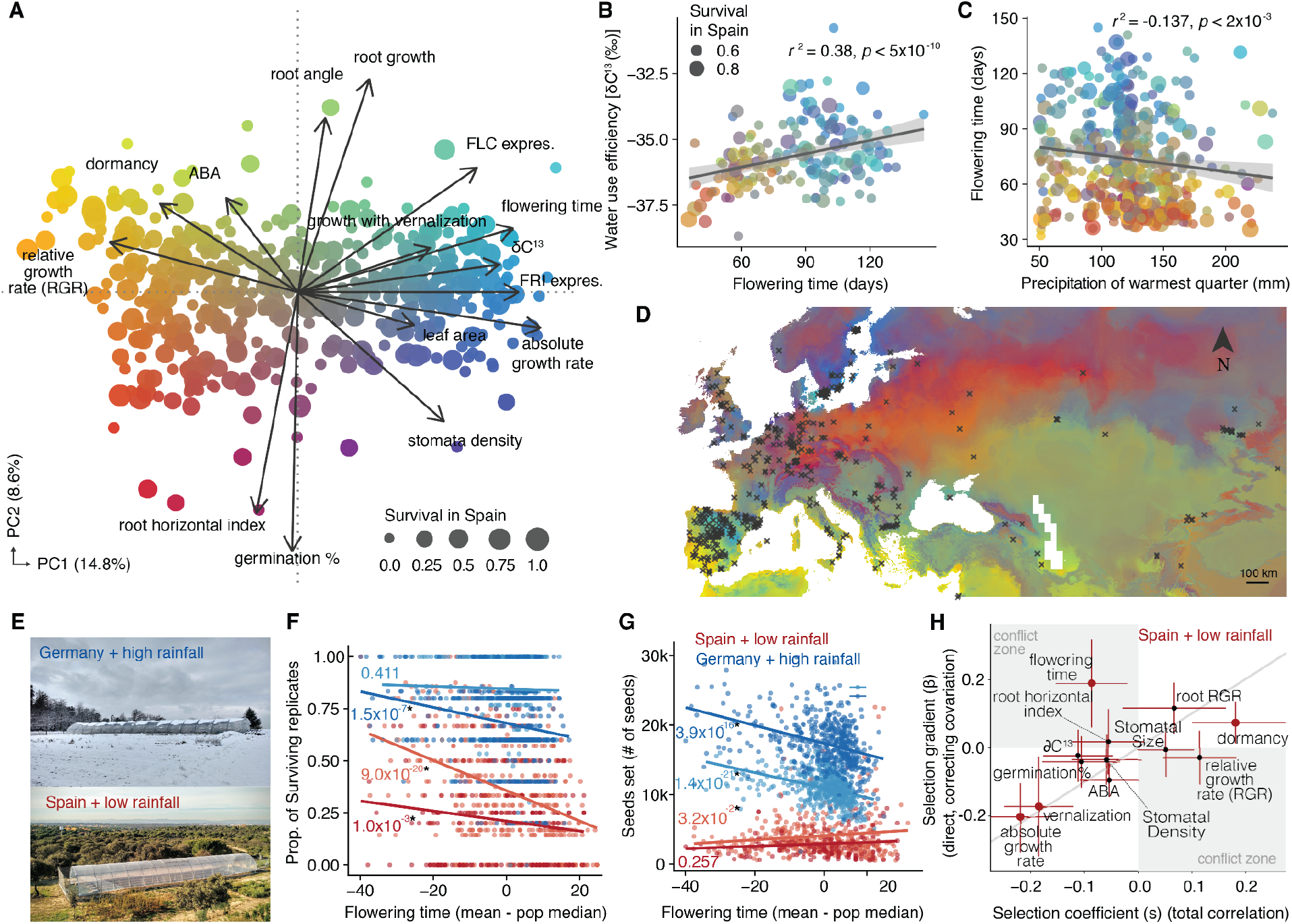
Escape and avoidance strategies across the *A. thaliana* native range leads to natural selection conflict. Principal Component Analysis (PCA) of 64 ecophysiological and life history traits in 515 accessions (arrows indicate loadings of key traits). Circle size indicates fitness in a Spanish field experiment ^33^ and a color gradient is mapped on the points to distinguish phenotype groups. (**Fig. S5**). Correlations between flowering time at 16°C and WUE (using δC^13^ as proxy) for 160/515 accessions, or (**C**) precipitation of the warmest quarter for 502/515 accessions (**Text SII**); colors correspond to those in (**A**). **(D)** Predicted phenotypes in color scale from (**A**) across the Eurasian native range of *A. thaliana* (**Text SII**). **(E)** Common garden sites. The absolute correlation between flowering time at 16°C (median-centered) and (**F**) survival as the proportion of surviving-to-reproduction replicates and **(G)** total number of seeds set ^33^. Associated *p* values (linear model) are in the same color as trend lines, with darker lines indicating low and lighter lines high planting density; note these are not selection coefficient *p* values but show the same patterns. (**H**) Total selection coefficients (*s*) vs. direct selection gradients (*β*, after accounting for trait covariation), using survival data from the hot and dry common garden experiment (Spanish field site), with low planting density. Large red circles indicate significant *β* and *s* (*p* < 0.05), small black circles indicate n.s.; error bars from 100 bootstrap replicate estimates (**Text SIII**); gray background areas indicate conflict zones where the signs (+/-) of total and direct selection conflict and are in the opposite direction.

To characterize what traits are adaptive in different climates, we leveraged *A. thaliana*’s broad native range, from Africa to Scandinavia, spanning extreme climate gradients from season lengths to water availability in the soil. High annual temperature is typically associated with early flowering and thus *escape* (Pearson’s *r*_flowering-mean,max.temp._ = -0.15, *p* < 5×10^-4^; **Fig. S2)**, while low precipitation during the growing season is associated with late flowering and thus *avoidance* (Pearson’s *r*_*ft-bio18*_ = -0.14, *p* < 1.5×10^-5^; **Fig. 1C**). These associations appear contradictory since high temperatures often coincide with low precipitation or water availability, and warming and drying are both expected to increase with climate change ^7^. Transplant experiments are the gold standard to understand adaptation by directly comparing the fitness of different populations and their phenotypes in specific environments. Such experiments often report natural selection favoring early flowering, low WUE genotypes ^3,20,26–28^ (**Table S9**), but in warm and dry climates such as Southern Spain ^29^, Eastern Spain ^26^, and Italy ^3^, flowering and fitness are sometimes uncorrelated, and in rare cases is somewhat paradoxically correlated with late flowering. Even in the same location, different precipitation patterns in different years can either favor early- or late-flowering ^27^, suggesting a complex relationship of climate with *escape* and *avoidance* strategies.

Mapping PC axes across the native geographic range using climate variables and a random forest model, we recapitulate the phenotypic landscape across a geographic landscape, where the early and late flowering phenotypes are represented at the extreme latitudes (**Fig. 1D**). At low and mid-range latitudes with short, mild, and wet winters, populations have evolved to not require vernalization to flower^30,31^ , and subsequently flower early ^28,32^, and rapidly complete their life cycle, canonical of the *escape* strategy to take advantage of a short rainy season. At the high latitudinal extreme of Scandinavia, populations live in climates with many months of snow cover, germinate in the fall, overwinter as small rosettes, and then have relatively low precipitation during key months of the life cycle from January to May (Pearson’s *r* = -0.15 *p* < 5×10^-2^; **Fig. S2**) and flowerlate in the season. Likewise, high elevation populations have characteristics similar to those of the northern latitudes.

To specifically estimate what trait combinations are adaptive in precipitation-limited conditions, we combined fitness data from rainfall-manipulated outdoor common garden experiments in Spain and Germany ^33^ with our imputed trait database and decomposed phenotypic space (**Fig. 1E**). The well-known observation in annual plants of a selective pressure over early flowering time is immediately apparent (*s*_*fitness*_ = -0.191, *p*_*boot*_ < 1×10^-3^; **Table S7**). Partitioning fitness data into survival to reproduction and fecundity, we observed the first incongruence of selection under the hot and dry Spain environment, where there was a significant advantage of early flowering for survival (**Fig. 1F)** (*s*_*survival*_ = -0.282, *p*_*boot*_ < 1×10^-3^; **Table S7**), but an advantage of late flowering for fecundity (**Fig. 1G)** (*s*_*fecundity*_ = 0.085, *p*_*boot*_ < 5×10^-2^; **Table S7**). Flowering time, together with other traits, is a main driver of phenotypic variation in PC1, which also exhibits an opposite direction of selection between survival and fecundity (*s*_*survival*_ = -0.239, *p*_*boot*_ < 1×10^-3^; *s*_*fecundity*_ = 0.109, *p*_*boot*_ < 5×10^-2^, **Table S6**), as do growth rate (**Table S9**) and primary seed dormancy (**Table S10**). This is consistent with delayed flowering permitting plants to accumulate more resources for reproduction but at an increased risk of mortality ^34^, however it could also indicate there are indirect selective pressures for flowering time influencing trait evolution ^35^.

Because PC1 includes several key traits underlying the trade-off between *escape* and *avoidance*, we wanted to disentangle phenotypic selection pressures by accounting for correlated traits in a multivariate selection analysis ^36^, only quantifying *direct* effects of natural selection in each of 12 key focal traits (**Fig. S3)**. Most traits remained consistent in their direct, correlation-free selection estimates. For instance, for absolute growth rate, the total selection estimate (*s*_*survival*_ = -0.219, *p*_*boot*_ < 10^-3^) was very similar to the direct selection estimate (β_*survival*_ = -0.206, *p*_*boot*_ < 10^-3^; **Fig. 1H**), indicating that there is no influence of correlated trait variation on selection for absolute growth rate. Likewise, the vernalization requirement for flowering time and primary dormancy selection estimates were also consistent (**Fig. 1H**). All of this evidence indicates selection for *escape* phenotypes. Strikingly, the sign of selection for flowering time became reversed from negative to positive, suggesting an advantage of late flowering accessions (*avoider*s) once we had accounted for trait correlations (*s*_*survival*_ = -0.087, *p*_*boot*_ < 10^-2^; β_*survival*_ = 0.190, *p*_*boot*_ < 10^-3^; **Fig. 1H; Table S13**). This strong positive estimate of direct selection indicates that there is a clear fitness advantage to late flowering. Likewise, this switch in the sign of selection estimates when accounting for correlated trait variation, particularly with growth rate, suggests this variation is having a strong impact on the selection pressure that flowering time experiences (leave-one-out re-analyses indicate this reversal can be seen when accounting for growth rate and vernalization see **Text SIII**).

A fitness advantage to late-flowering accessions of *A. thaliana* in hot and dry environments is not completely unexpected, as these plants are more resilient to water limitation ^20,37^, often manifested by water accumulation through alterations in root development, stomata control, and overall WUE. Similar to flowering time but to a lesser extent, we also observed a shift in the direction of selection in the WUE proxy δC^13^, where total selection was negative (*s*_*survival*_ = -0.112, *p*_*boot*_ <10^-3^) but the direct effect was non-significant (**Fig. 1H; Table S13**). While we expected phenotypic constraints from these correlations to be pervasive across species, direct selection being so strongly inconsistent with total selection is a rare observation relative to many plant and animal species and thousands of measured morphological, phenological, or physiological selection estimates ^35,38^. However, given flowering time is such a fundamental trait in trade-off with other water use traits, this conflict of selection will likely have broad implications in impeding multivariate trait evolution in plants.

Does this selection conflict at the trait level then also manifest at the genetic level ^4,39,40^? To answer this question, we performed systematic genome-wide association (GWA) analyses with the accessions from the 1001 Genomes Project ^24^ for each of the 12 key traits ^41^, including pairwise multivariate linear mixed model (mvLMM) GWA ^42^ to estimate genetic correlations and shared genetic architecture. Most traits had moderate-to-high SNP-based heritability *h*^*2*^ = 0.55 [0.25-0.85]) and were polygenic (*γn* = 48.5 causal loci [21.6-75.4]) (**Table S15**). The strongest, most consistent genetic correlation was between flowering time and growth rate (*r*_*g-flowering-growth*_= 0.45, 95% CI 0.38–0.52; **Fig. S17**), and both were in turn strongly correlated with WUE at the genetic level (*r*_*g-flowering-WUE*_= 0.34, 95% CI: 0.26–0.42, **Fig. 2A**; *r*_*g-growth-WUE*_= 0.33, 95% CI 0.25–0.42; **Fig. S17**). To further investigate the effects of specific genomic loci on flowering time and WUE and their signatures of natural selection, we examined the effect sizes on both traits of the top 0.05% SNPs (n=274) in our mvLMM GWA. This showed a striking parallel; alleles increasing WUE also increased flowering time, and vice versa (Fisher’s Exact Test odds ratio 2.7×10^-4^, *p* < 2.2×10^-16^; **Fig. 2B**). The expected allele frequency changes (**Fig. 2C**) show a static trend where putatively pleiotropic loci impacting both traits sometimes increase and sometimes decrease in frequency implying evolutionary stasis. To solidify this inference, we applied the Breeder’s equation, which accounts for genetic correlations amongst traits and heritability to predict the average phenotype change of each trait (*sensu* Lande & Arnold, ^*36*^; **Fig. 2D**). Only traits with consistent total (*s*) and direct (*b*) selection estimates, as well as high heritability, such as growth rate, were predicted to have a significant change in the mean trait value in the next generation (**Fig. 1H**). Consistent with the constraints of selection on flowering time and WUE, and the low heritability of the latter, we found no significant shift in the mean trait value. Rather, we predicted the counter-intuitive “evolutionary stasis” of these traits in a Mediterranean climate with water limitation (Δ*z*_flowering_=0.136, p-value > 0.05; Δ*z* _WUE_=-0.001, p-value > 0.05, **Fig. 2D, Table S17**). In conclusion, though the estimate for flowering time is leaning positive, the influence from correlated trait variation and selection on growth rate is constraining the evolution of late flowering time.

**Fig. 2.**
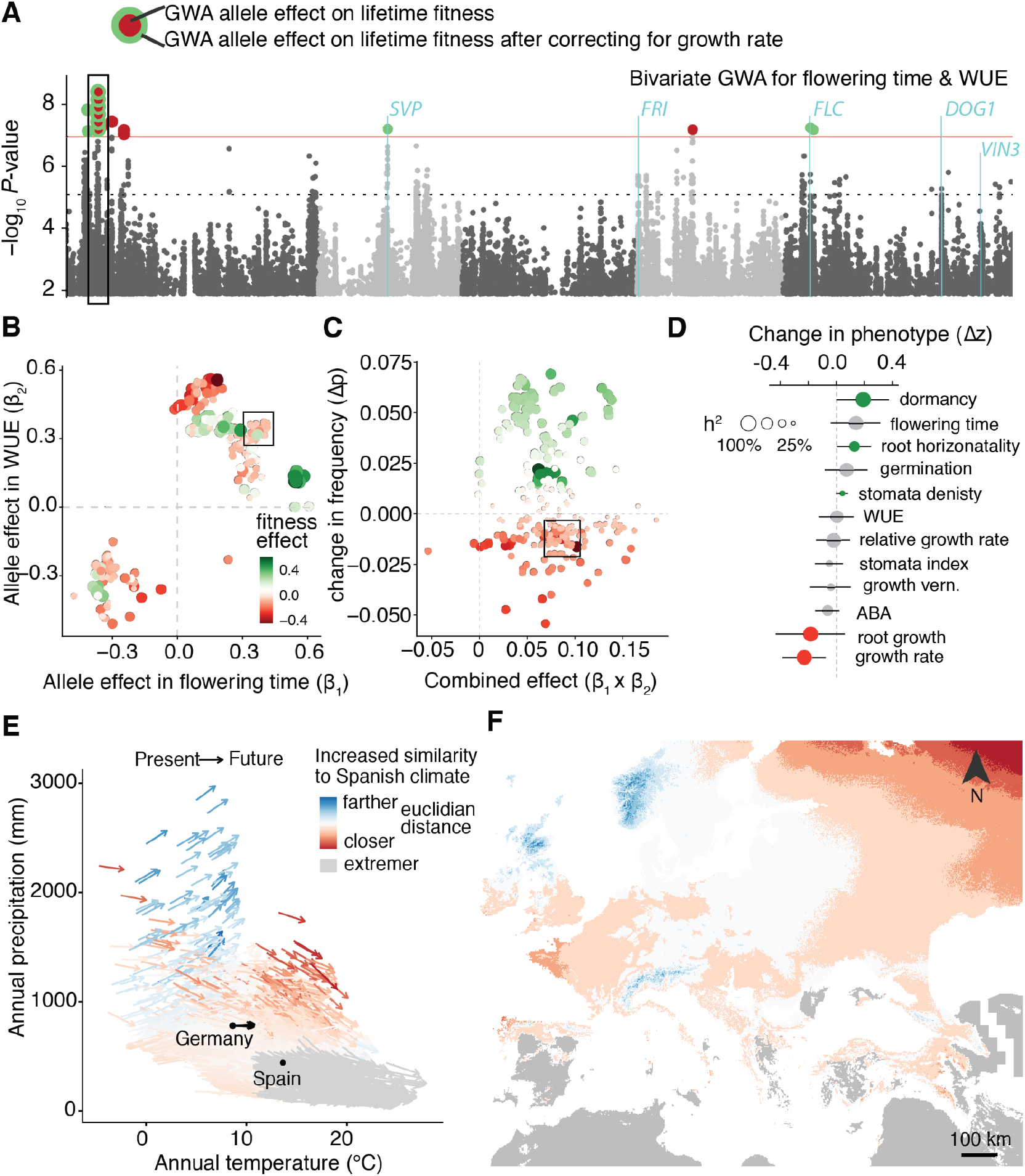
The joint genetic architecture of *escape* and *avoidance* constrains trait evolution in water-limiting environments. Manhattan plot of mvLMM GWA of flowering time and WUE (δC^13^) (n=248). Red line is the Bonferroni-corrected genome-wide significance threshold of -log_10_ *P-*value > 7.03, black dashed line indicates the top 0.05% of alleles with -log_10_ *P-*value > 5.13; annotated genes are significant for flowering time ^15^ and are identified in our extended analysis of these data (**Text SIV**). Significantly associated SNPs are colored based on the effect size of a separate GWA with lifetime fitness from the hot, dry Spanish environment, where allelic effects on fitness were estimated; circle color indicates alleles with a positive (green) or negative (red) association with fitness; outer circle color of alleles on chromosome 1 indicate the effect each allele has on fitness after correcting for growth rate. **(B)** Top 0.05% SNP effect sizes for flowering time (ß_1_) and WUE (ß_2_) from GWA described in **A**, circles colored and sized by lifetime fitness effects estimated from fitness GWA described above. (**C**) Expected allele frequency change in response to selection (**Text SIV**) by the combined allelic effects for flowering time and WUE. (**D**) Predicted mean phenotypic response to selection across traits, size of circles indicates SNP-based narrow-sense heritability estimate represented as a %, color of the circles indicates positive (green), negative (red), or no (gray) predicted change in the mean trait value of the next generation. (**E**) Predicted change in annual precipitation and temperature similarity to the Spanish field site climate in 2050 (measured as Euclidean distances). (**F**) Map of climate trajectories in **A**. Climate maps were retrieved from Max Planck CMIP5 IPCC projections of business-as-usual scenarios (worldclim.org). Gray areas indicate regions where temperature will increase while precipitation will decrease below the average precipitation at the Spanish field site (*i*.*e*. more extreme conditions).

We therefore wondered whether any major effect alleles for flowering time and WUE were under the same natural selection conflict that was pervasive at the trait level. Many of the major effect alleles are in well-known flowering-time genes, including *FLC* and *SVP* **(Fig. 2A**). In addition, an imputed GWA identified other flowering-time genes *FRI*, known to have pleiotropic effects in flowering time and WUE^20^, *DOG1*, and *VIN3* ^*15*^ as having major effects (**Fig. S15**). We also discovered a highly significant locus on chromosome 1 as appearing consistently in our different analyses (**Text SIV**). The top SNPs span a region of ∼7.5 kb and 5 candidate genes (**Fig. S18; Text SIV)**, with the expression of three of these genes correlating with flowering time ^43^, and another with WUE (**Fig. S19**). To estimate the effect of natural selection on these alleles, we conducted an additional GWA with fitness from Spain hot/dry conditions ^33^. The chromosome 1 alleles that increased both flowering time and WUE (**Fig. 2B**) were negatively correlated with fitness (mean effect size = -0.054 [0.10]; **Fig. 2A**). After including growth rate as cofactor though, their correlation with fitness reversed (mean effect size = 0.173 [0.190]; **Fig. 2A**), meaning after the negative selective effect from growth rate was accounted for, the remaining variation in fitness suggests positive selection for these alleles. These same alleles also correlated negatively with survival (mean effect size= -0.117 [0.089]) but positively with fecundity (mean effect size= 0.089[0.070]). We speculate that this locus is pleiotropic and subject to antagonistic selection, recapitulating the selection constraint originally observed at the trait level between flowering time and growth rate.

That two opposing adaptive traits controlled by partially-overlapping gene variants are selected under warm and drought conditions in *A. thaliana* does not bode well, as it is expected that the climate in much of Europe will become more similar to the Spanish hot/dry environment, potentially increasing the natural selection conflict in these traits, exacerbating this evolutionary trade off (**Fig. 2E, F**). We experimentally tested this hypothesis by reducing rainfall in Germany, simulating Spanish rainfall. Compared to the well-watered treatment where early flowering time is negatively selected (β_*survival*_ = -0.022, *p*_*boot*_ < 5^-2^; **Table S14)**, reduced rainfall in Germany led to flowering time being less important for fitness and WUE (which was non-significant in well-watered conditions) becoming positively correlated with survival (β_*survival*_ = 0.067, *p*_*boot*_ < 10^-3^; **Fig. S7**). This is direct evidence that a German climate becoming more similar to a Spanish climate would lead to the same natural selection conflict. Given that the onset of flowering in many species is accelerating ^8,44,45^, combined with the universality of escape and avoidance strategies across plant clades ^22^, it is of utmost relevance to understand whether trait constraints could be released in order for optimal phenotype combinations to be able to evolve.

In such a constraint-of-selection scenario, we may expect that only large-effect alleles for highly heritable traits may be efficiently selected and become fixed ^36^. To test this expectation directly, we simulated the scenario of a large-effect allele accelerating flowering time to become fixed in a population through CRISPR/Cas9 engineering. We asked how this would impact trait correlations with WUE and whether it would release indirect natural selection constraints, permitting evolution of WUE traits. We used *FLOWERING LOCUS C* (*FLC*) as a proof-of-principle large-effect gene, because *FLC* encodes a master regulator that prevents premature flowering in winter. Our GWA analyses had identified natural *FLC* alleles as controlling the bulk of flowering time variation (**Fig. S14**), and the mvLMM GWA suggests *FLC* may play an important role in the shared variation between flowering time and WUE (**Fig. 2A**). In natural populations, *flc*-knock-out (KO) and severe knock-down (KD) mutations nearly always result in an early spring-flowering phenotype and segregate in warm environments at the Southern extent of the range ^31,46^. It is therefore theoretically possible that a *flc*-KO mutation could become fixed in warm environment populations and accelerate flowering time for an entire population.

We designed gRNAs in two downstream and two upstream target regions of *FLC* (**Fig. 3A**) across a diverse set of 62 accessions. Overall, 84% of the edited lines had reduced *FLC* transcript levels, with 39 likely having large deletions (KOs) and the remainder having weaker mutations (KDs) (**Fig. 3A**). As flowering time is correlated with WUE, we expect modulation of *FLC* activity to have a pleiotropic effect on WUE. We measured flowering time and δC^13^ for a subset of the *flc* lines and their wild-type counterparts (n=19), along with Col-0 wild type (**Fig. 3B**). In most *flc* lines, flowering was accelerated (mean [se] = 42.61 days [0.10] compared to wild-type of 106.54 days [0.36]) (**Fig. 3B**). There was also a significant reduction of WUE (*t* test, Benjamin-Hochberg correction, *p*_*adj*._ < 0.05; **Fig. 3B**). Remarkably, the relationship between flowering time and WUE was 2.7 times weaker amongst the *flc* lines compared to the wild types (linear regression flowering ∼ βwue; β_*flc*_ [se] = 4.58 [2.2], β_*wt*_ [se] = 10.44 [16.9]), clear evidence for a relaxation of phenotypic genetic constraints in the *flc* population (**Fig. 3B**), and theoretically enabling natural selection to act on WUE.

**Fig. 3.**
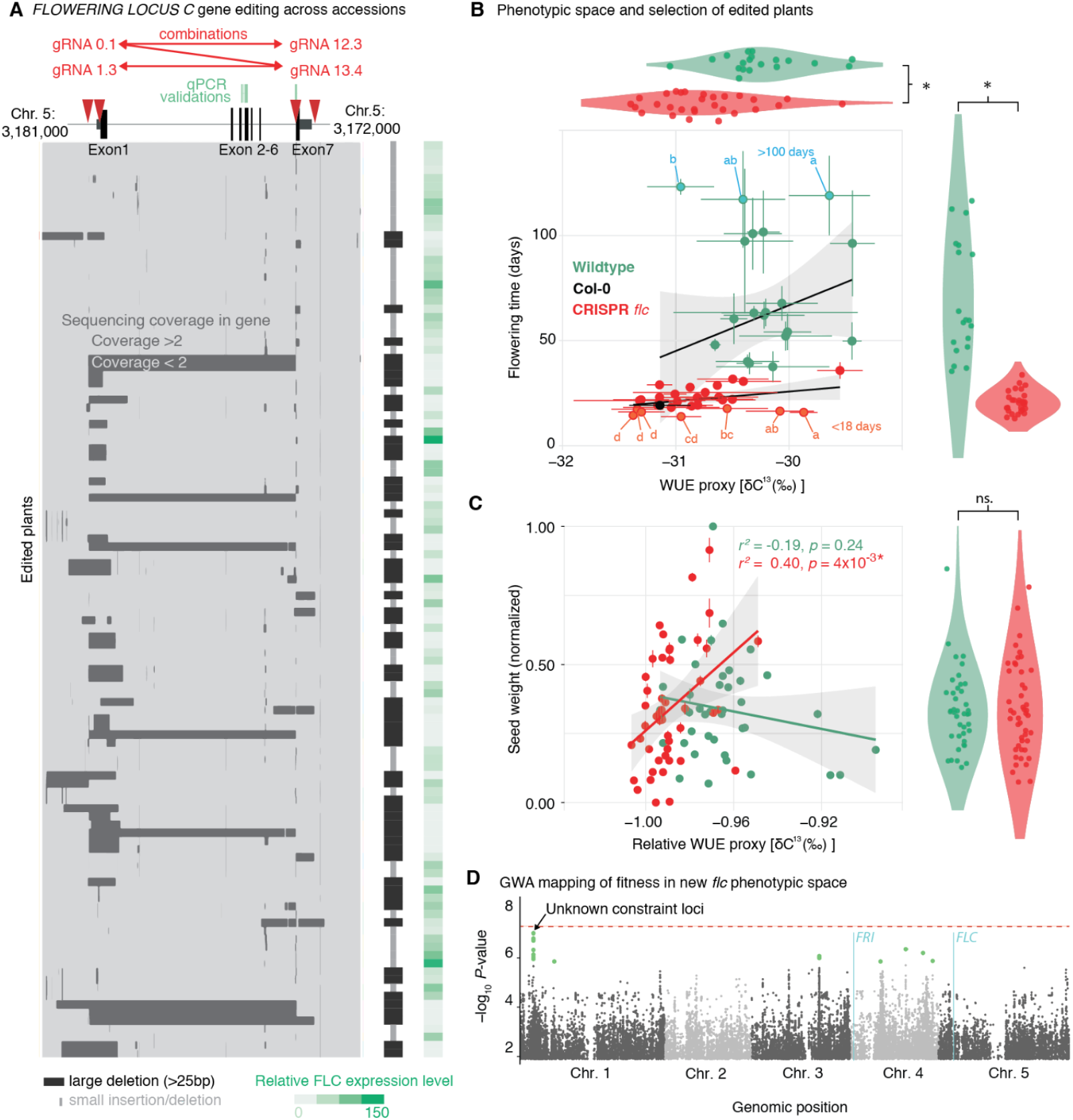
CRISPR/Cas-mediated gene editing of diverse *A. thaliana* accessions relaxes trait correlations between flowering and WUE that can become unfavorable in future. (**A**) Top, diagram of the *FLC* locus. Black boxes represent exons 1 - 7, gRNA target sites indicated by red arrows. The red connectors indicate gRNA combinations in single editing constructs. Bottom, whole-genome sequencing coverage at the *FLC* locus in different lines. Dark gray signifies reduced coverage indicative of deletions. Each row represents one *flc* mutant line. On the right, membership of each line in a “large deletion” and “small mutation” class is indicated. On the far right, relative *FLC* expression in nine-day-old plants grown in long days is shown. (**B**) Correlation of flowering time and δ^13^C for *flc* mutants and wild types in 22°C under LD. Red indicates *flc* mutants and blue wild types. Different letters indicate significant differences in δ^13^C (ANOVA with post hoc Tukey HSD, *p*_*adj*_ < 0.05). (**C**) Correlation of seed weight and WUE for *flc* mutants and wild types in warm conditions without water limitation. Relative WUE is δ^13^C compared to δ^13^C of the Col-0 accession. (**D**) Manhattan plot of mvLMM GWA of flowering time and relative WUE for the *flc* lines, with points colored by selective effect on alleles measured by univariate GWA using fitness of *flc* lines in warm conditions without water limitation.

To test whether natural selection on flowering time and WUE would change in our *flc* population, we grew these lines (n=50) in warm conditions without water limitations (21°C under long days (LD, 16 h light / 8 h dark). Quantifying fitness as seed yield, we now found a positive relationship of fitness with WUE in the *flc* lines while we still found the canonical constrained negative fitness relationship with WUE in the corresponding wild-type population (**Fig. 3C**). Some accessions even had the unique combination of early flowering and high WUE phenotypes. These results are significantly different from a longstanding observation in annual plants like *A. thaliana*, where phenotypic selection for WUE is most elusive ^20^. We speculate that it may be the strong effect of life history strategy in fitness that may have hindered the observation that high-WUE accessions do in fact perform better under hot, high evapotranspiration conditions. We finally performed a mvLMM GWA on these 50 *flc* mutant lines using the flowering time and WUE trait data and assessed the selective effects on alleles using the fitness data from the experiment. We found that the alleles on chromosome 1 that contribute to increased WUE (**Fig. 3D**) are now under positive selection (mean effect size = 0.0082 [0.002]). These results support the idea that some large-effect loci, despite their pleiotropic effects, may change the landscape of multi-trait correlations enabling the evolution of novel trait combinations beyond what we had theoretically predicted.

Future climate change is expected to generate novel environments to which species have not yet been exposed in the past and may thus not have evolved the optimal phenotypic combinations. Using gene editing, we have simulated a possible future population that had a large-effect loss of function mutation become fixed, drastically impacting phenotype distributions and the direction of selection. Phenotypic variation within *A. thaliana*, and likely other species, is shaped by genetic variation and past selection forces that have generated an *escape*-*avoidance* strategy continuum. Given that rapid climate change has already led to early flowering in many species ^8–10^, we expect natural selection to act along “axes of least resistance”, that is the most easily accessible portions of phenotypic space due to high heritability and low polygenicity, even if other beneficial strategies are theoretically possible. Although predicting future evolutionary trajectories is complex, we speculate that large-effect alleles affecting flowering time exist in natural populations, and that these will rapidly increase in frequency due to natural selection in a hotter-and-dryer native climate. After the first response to selection, we predict increased WUE will be selected for. However, the short-term evolution of early flowering time will have a detrimental pleiotropic effect in WUE, the consequences of which for adaptation are so far unknown. The expanding genomic and molecular biology knowledge across plant species and democratization of gene editing technologies should enable us to disentangle complex trait correlations, which is an underestimated constraint to evolution that could lead species to local minima in the fitness landscape that are difficult to escape.

## Materials and Methods

### Trait data curation

The database of 1,862 traits was assembled from 108 published sources encompassing a combination of laboratory and field experiments that use a subset of *A. thaliana* accessions from the 1001 Genomes Project (**Table S1; Text SI**). The trait data are averages for a genotype across replicates measured in replicated environments and were not measured in a single experiment or for single plants. We classified traits into a general functional category, with 1,282 traits as being related to one of three drought adaptation strategies; *escape* (n=127), *avoidance* (n=382), and *tolerance* (n=773) (**Fig. S1**). Because most traits had only been measured on subsets of1001 Genomes accessions, with subsets overlapping to different degrees (**Fig. S1**), we used the R package *missForest* to fully impute the trait dataset (see **Text SI; Table S2**).

### Trait associations

For PCA, traits related to seed dormancy, vernalization, germination, flowering time, leaf traits, roots, stomata, growth rate, and stress response were isolated (n=205). Highly correlated traits were removed (*R*^*2*^ > 0.7), resulting in a total of 64 usable traits (**Fig. S1)**. We estimated how many of the 1,862 traits were correlated with four flowering time datasets, WUE, and growth rate, using Pearson’s correlation coefficient with a significance threshold of 0.05 and the non-imputed trait dataset (**Table S3**). We also downloaded bioclimatic, temperature, precipitation, and evapotranspiration rate estimates from WORLDCLIM 2.0 for all localities associated with the 515 accessions for which fitness data were available, and estimated Pearson’s correlation coefficient between climate data and these six target traits (see **Text SII; Table S4**).

### Selection estimates

We computed total selection coefficients *s* (Lande and Arnold 1983) for all non-fitness traits (**Table S5**) and for 20 PC axes explaining 75% of trait variation (**Table S6)**, highlighting focal traits such as flowering time (**Tables S7, S8**), growth rate (**Table S9**), seed dormancy (**Table S10**), and WUE (**Table S11**). Total selection was calculated as the covariation between relative lifetime fitness *w* and a given trait *z* (mean and variance centered): *w* = *sz*, where *s* is the total selection coefficient. Fitness data were from an outdoor common garden experiment in two different environments, Madrid, Spain (m) and Tübingen, Germany (t) (**Figure 1E**) ^33^. Fitness data were mean-centered, and trait data were mean-centered and variance scaled prior to estimating *s*. Additionally, *s* was estimated from 100 bootstrap replicates with replacement in order to generate confidence intervals and determine significance.

To understand the direct effects of natural selection on traits, we accounted for correlations amongst traits using the multivariate selection approach, β = *P*^−1^*s* , where *P* is the phenotypic variance-covariance matrix ^36^, *s* is the previously quantified total selection coefficient (without accounting for correlations), and β is the gradient of selection (or direct selection) that captures the independent effects of each trait that cannot be attributed to their associations with other traits. Through a series of trait decorrelation, and expert selection, we narrowed the analyses to 12 focal traits related to both seasonal and drought adaptation (**Table S12, Fig. S3**). We focused this analysis by only using fitness data from the low water Spanish site (hot and dry; **Fig. 1H, Table S13**) and the high water German site (cool and wet; **Fig. S4, Table S14**) (see **Text SIII)**.

### Genome wide associations

To perform genome wide association (GWA) analyses, we started with the 1,135 accessions and 11,769,920 SNPs from the 1001 Genomes Project (1001 Genomes Consortium 2016). Depending on trait data coverage, not all samples and SNPs were used as we ran GWA on first the focal 12 traits, but also on all traits (n=1,862) using the raw data, quantile-normalized data, and imputed data. For each trait we conducted GWA using a univariate linear mixed model (LMM) implemented in GEMMA (ref. ^41^). This model corrects for genome-wide background differentiation collinear to the phenotype of study (i.e. relatedness and population structure correction). No minor allele frequency (MAF) cutoff was used unless stated (**Text SV**). Additionally, using the original imputed SNP matrix from http://arapheno.1001genomes.org, which contains 10,709,466 SNPs, we used a subset of 1,353,386 SNPs for all 1,135 1001 Genomes accessionsin a Bayesian sparse linear mixed model (BSLMM) GWA, also implemented in GEMMA (Zhou and Stephens 2014), for each of the 1,862 phenotypes (see **Text SVI**). For the 12 target traits used in the selection analysis (**Table S16**), pairwise multivariate LMM (mvLMM) GWA were performed ^42^. Again, analyses were run with non-imputed and imputed data, and when the data were not imputed only 16/66 pairs of mvLMM GWA were possible given lack of overlapping samples and large number of parameters to estimate (**Text SIV**).

### Fitness GWA

Fitness data from field experiments of 515 accessions ^33^ were used in a univariate LMM GWA with no MAF cutoff and corrected for population structure using a kinship matrix. The sign of the effect size estimates for each allele indicated whether the allele has a positive or negative effect on the quantitative fitness measure. We use these effect size estimates to color alleles to indicate positive (green) and negative (red) selection (**Fig. 2A, B, C**). To account for the variation in fitness that is explained by absolute growth rate variation, we estimated a linear model of fitness explained by growth rate and then used the residuals from this model in an additional LMM GWA to see the remaining selective effect on alleles (**Fig. 2A**).

### Prediction of phenotypic response to selection

We inferred the population responses to this natural selection by using the Breeder’s Equation or selection response equation (ref. ^49^), Δ*z* = *GP* ^−1^ *s*, where *G* is the additive genetic variance-covariance matrix constructed from heritability and genetic correlation estimates among 12 traits, *P* is the phenotypic variance-covariance matrix, and *s* is the raw covariance of the trait with lifetime fitness in hot-dry conditions. For this, we used the estimates of *s* from lifetime fitness in all environments from ref. ^33^, but primarily report the hot-dry environment (Madrid (m) and low water (l)), but also mention results for WUE in the cool-dry environment (Tübingen (t) and low water (l)). For the genetic variance-covariance matrix, we used the SNP-based heritability estimates of target traits using the GWA results (**Table S16**), and the genetic correlations estimated from the mvLMM GWA from imputed data (**Fig. S18**).

### Climate projections

We downloaded the averaged historical (1970-2000) bioclimatic variables ^47^ for the Eurasian range of *A. thaliana* at 5 km^2^ (2.5 minutes) resolution. Future predicted climate data were retrieved from the Max Planck CMPI5 IPCC model of a business-as-usual scenario ^47^. We plotted the 1970-2000 average value of annual precipitation and mean annual temperature as the beginning of the arrows (**Fig. 2E**) and the 2050 predicted values as the end of the arrows to indicate the trajectory of the climate in precipitation and temperature space. Arrows indicate whether the euclidean distance (of both temperature and precipitation) between the current and 2050 climates becomes more similar to the current Spanish climate (red) or more dissimilar (blue) (**Fig. 2E**). Trajectories with predicted precipitation and temperature more extreme than the Spanish site are in gray. The intensity of the trajectory was then mapped back to the initial localities across the Eurasian range to visualize where in the native range it will become more like the Spanish or German climate (**Fig. 2F**).

### CRISPR/Cas9 gene editing of FLC

We selected 60 late- and two early-flowering accessions of *A. thaliana* from the 1001 Genomes Project ^24^, that represent 13 out of 25 of the major FLC haplotypes (**Fig. S21B** and **Table S23**), for CRISPR/Cas9 gene editing. gRNAs were designed using CCTop and no off-target binding was predicted ^48^. For CRISPR/Cas9 mutagenesis, we used a slightly modified previously established toolset ^49^ with a supermodule destination binary vector carrying a plant-codon optimized Cas9 driven by a *UBQ10* promoter (**Text SVI**). We additionally mutated a TTTT stretch to TTCT by introducing a unique XbaI site. Constructs expressing pairs of four gRNAs (0.1 and 13.4, 1.3 and 13.4, 12.3 and 0.1; **Fig. S21D**) were transformed as pools into the *Agrobacterium tumefaciens* strain ASE ^50^. Accessions were transformed with the floral dip method ^51^ (**Text SVI**).

### Mutant screen

T_1_ seeds from all pots per accession were pooled and screened for seed fluorescence or absence thereof (**Text SVII**). Up to 15 positive primary transformant seeds (T_1_) were sown and treated with a heat cycle to increase Cas9 editing efficiency according to ^52^. Seeds from all T_1_ plants that reached the generative phase were bulked per accession to generate T_2_ seed pools.

From the T_2_ seed pools, early flowering individuals were selected by screening more than 1000 plants per accession. Approximately 25 early flowering plants per accession were propagated to the T_3_ generation (totaling more than 2,500 plants from more than 120,000 screened individuals). T_3_ seeds were examined for presence of the transgene using the fluorescence reporter system. Flowering time was analyzed for one to six T_3_ lines (five plants each) alongside five wild-type plants. One plant from one to two lines per accession of early flowering, non-segregating lines was selected for sequencing using Illumina 2x150 bp reads (see **Text SVII**).

### Expression analysis

Three biological replicates per wildtype and one or two mutants were randomly distributed over three plates, grown at 22°C under long days (LD, 16 h light / 8 h dark), and sampled at ZT14 (two hours before dark). Several seedlings per line were pooled as one biological replicate. RNA from each biological replicate was extracted using a column-based protocol ^53^. 1 μg total RNA was reverse transcribed with RevertAid First Strand cDNA Synthesis Kit (Thermo Fisher Scientific, Waltham, USA) using an oligo (dT) primer and cDNA was analyzed in a CFX384 Real-Time System Cycler (BioRad, München, Germany). The relative quantification was calculated with the ΔΔCt method using *ACT8* (AT1G49240) as a standard ^54^ and calibrated by biological replicate 1 (rep1 ) of Col-0.

### Measurement of carbon isotope ratio

Mutant and wild types contrasts were grown in three pots representing three biological replicates at 22°C under LD. Rosettes were harvested at either the initiation of flowering or after 22 days after germination and tissue was dried at 60°C for 24 hours and homogenized. Dried material for 19 mutants and wild types was sent to Isolab GmbH (Schweitenkirchen, Germany) for an analysis of carbon isotope composition (δ^13^C) with ^13^ C-CF-IRMS. Four technical replicates per sample were analyzed. For a more detailed description of the procedure, see ^55^. Data are presented as δ^13^C [‰] vs. V-PDB (**Fig. S25C**).

### Selection experiment with flc mutants

50 mutant / wild-type contrasts, including a biological replicate, germinated and were grown in warm, non-water-limiting conditions in a growth chamber at 21°C LD. Plants were bottom watered 2 times a week. Flowering time was measured as days to first petal emergence and seed weight was measured as total weight of seeds harvested (mg). Values for flowering time and fitness were averaged across replicates. Given the conditions were comparable, instead of re-measuring δ^13^C for the 50 wild-type accessions again, we assumed ranks of δ^13^C across accessions are relatively constant across environments based on previous research ^20^, and imputed the data based on a larger δ^13^C dataset of several hundred accessions ^25^ (NRMSE = 0.13; **Table S2**). For the *flc* mutants, we used the 19 values of δ^13^C measured above, and then imputed the values for the remaining 31 accessions using those 19 values, the δ^13^C data from ^25^ (**Fig. S26**), and the flowering time of the accessions from this experiment in order to ground the values predicted to this experiment and not general values of δ^13^C. Likewise, for both the wild-type and mutant δ^13^C data, we scaled the values using δ^13^C values from Col-0 grown in similar conditions, thus making the measure a relative measure of WUE (see **Text SVIII**).

## Data availability

The 1,862 phenotypes imputed for 1,135 wild *Arabidopsis thaliana* strains are available at github.com/moiexpositoalonsolab/Athaliana_Natural_Selection_Conflict/data. The data from the 1001 Genomes of *Arabidopsis thaliana* are available at 1001genomes.org. The seed collection can be obtained at the Arabidopsis Biological Resource Center (ABRC) under the ID CS78942. The code to reproduce analyses and figures is available at github.com/moiexpositoalonsolab/Athaliana_Natural_Selection_Conflict. Reads were deposited at NCBI SRA with accession number: <TBD upon publication>. Seeds were deposited at NASC with the accession number <TBD upon publication>.

## Acknowledgements

We thank Y. Voichek for their effort in the collection of trait data. We thank the Exposito-Alonso, Rhee, and Weigel Lab members, and Oliver Bossdorf for comments and discussion. We thank S. Schäfer, N. Betz, A. Spazierer, F. Strauss, N. Vasilenko, and J. Kreienbrink for technical assistance in genome editing. This work is supported by the Office of the Director of the National Institutes of Health’s Early Investigator Award with award # 1DP5OD029506-01 (M.E.-A.); by the U.S. Department of Energy, Office of Biological and Environmental Research, grant # DE-SC0021286 (M.E.-A., S.Y.R.), DE-SC0018277 (S.Y.R.), DE-SC0008769 (S.Y.R., M.R.), DE-SC0023160 (S.Y.R.); by a USDA NIFA, grant # 2022-67019-36366; and by the U.S. National Science Foundation’s DBI grant # 2213983 (Water and Life Interface Institute (WALII)) (S.Y.R., M.E.-A.); Max Planck Society and Novozymes Prize of the Novo Nordisk Foundation (D.W.); by the Carnegie Institution for Science (M.E.-A., S.Y.R.); and the Howard Hughes Medical Institute and the University of California Berkeley (M.E.-A.). M.R. is supported by the National Science Foundation Plant Genome Project as a postdoctoral fellow (Grant No. 2109868). L.L. is supported by the National Science Foundation Graduate Research Fellowship (Fellow ID: 2020304890). S.H. is supported by Stanford’s Center for Computational, Evolutionary, and Human Genomics. For computational analyses, we relied on the High-Performance Computing clusters Calc and MoiNode supported by the Carnegie Institution for Science. This work was done in part on the ancestral land of the Muwekma Ohlone Tribe, which was and continues to be of great importance to the Ohlone people.

## Author contribution

M.E.-A., D.W., M.R., U.L., and S.Y.R. conceived the project. M.R., L.L., S.H., U.L., W.Y., and M.E.-A. conducted analyses, U.L. performed genome editing, U.L. and L.L. conducted experiments. All authors interpreted analyses and discussed results. M.R., U.L., D.W., and M.E.-A. wrote the manuscript with the input of all authors.

