## Supplemental Text and Figures for "Selection constraints of plant adaptation can be relaxed by gene editing"

### Gene editing relaxes pervasive antagonistic selection constraining plant adaptation

Supplemental Text: Extended text on analyses and results 5
I. Trait data 5
I.1 Data curation 5
I.2 Water Use Efficiency 5
I.3 Trait imputation 6
II. Trait analyses 6
II.1 Principal component analysis 6
II.2 PC climate models 6
II.3 Trait associations 7
II.4 Climate change projections 7
III. Selection estimation 8
III.1 Total selection (s) 8
III.2 Direct selection () 9
IV. Genome wide associations 10
IV.1 Linear mixed models 11
IV.2 Bayesian sparse linear mixed model 11
IV.3 Multivariate linear mixed models 12
IV.4 Genetic correlation approximations 13
IV. 5 Fitness effects 13
V. Prediction of phenotypic response to selection 13
VI. Gene editing of FLC 14
VI.1 Establishing lines 14
VI.2 CRISPR/Cas9 protocol 14
VI.3 Plant transformation 15
VII. Characterize FLC mutants 15
VII.1 Mutant screening 15
VII.2 Characterization of transgene insertions 16
VII.3 qRT expression analysis 17
VII.4 Measurement of carbon isotope ratio 18
VIII. Growth chamber experiment with FLC mutants and wildtypes 18
VIII.1 δ13C imputation 19
IX. Supplementary Tables 20
Supplementary Table 1. Summary of trait data assembled in this study. 20
Supplementary Table 2. Trait imputation accuracy. 20
Supplementary Table 3. Trait associations with flowering time, WUE, and growth rate. 20
Supplementary Table 4. Significant associations between target traits and climate variables. 20
Supplementary Table 5. Total selection coefficients estimated for all non-fitness traits. 20
Supplementary Table 6. Total selection coefficients estimated for PC axes 1 and 2. 20
Supplementary Table 7. Total selection coefficients estimates for all flowering time data. 20
Supplementary Table 8. Total selection estimates for flowering time traits from other studies. 20

Supplementary Table 9. Significant total selection estimates for growth rate related traits. 20
Supplementary Table 10. Significant total selection estimates measured for dormancy traits. 20
Supplementary Table 11. Significant total selection estimates for WUE (δC13). 20
Supplementary Table 12. Overlap of trait data with corresponding fitness data. 20
Supplementary Table 13. Total and direct selection estimates in the Madrid, low water site. 20
Supplementary Table 14. Total and direct selection estimates in the Germany, high water site. 20
Supplementary Table 15. Heritability and GWA parameter estimates for all traits. 20
Supplementary Table 16. Heritability and GWA parameter estimates for 12 key traits. 20
**X. Supplementary Figures** **21**
Supplementary Figure 1. Characterization of trait data. 21
Supplementary Figure 2. Climate associations for flowering time and WUE. 22
Supplementary Figure 3. Pairwise Pearson's correlation coefficients for 12 focal traits. 23
Supplementary Figure 4. Total selection coefficients vs. direct selection gradients. 24
Supplementary Figure 5. Total selection coefficients vs. direct selection gradients from the Germany, high water site. 25
Supplementary Figure 6. Total selection coefficients vs. direct selection gradients without growth rate and vernalization.Figure SII.4 Multivariate selection analysis for 10 focal traits (growth rate and vernalization growth removed) with dry-hot, cool-wet, and high/low planting density fitness data from Exposito-Alonso et al. 2019. 26
Supplementary Figure 7. Total selection coefficients vs. direct selection for low rainfall German site. 27
Supplementary Figure 9. Total selection coefficients vs. direct selection gradients for 26.Multivariate phenotype selection using fitness data from Fournier-Level et al. 2011 27
Supplementary Figure 10. Total selection coefficients vs. direct selection gradients for 27. 29
Supplementary Figure 11. Trait correlation data after imputation and normalization. 29
Supplementary Figure 12. Comparison of effect size estimates. 31
Supplementary Figure 13. Boxplot of heritability for avoidance and escape traits. 32
Supplementary Figure 14. Flowering time and WUE GWA. 33
Supplementary Figure 15. Flowering time and WUE mvLMM GWA. 34
Supplementary Figure 16. QQplots for flowering time and WUE mvLMM GWA. 35
Supplementary Figure 17. Flowering time and growth rate mvLMM GWA. 36
Supplementary Figure 18. Genetic correlation estimates between key traits. 37
Supplementary Figure 19. Flowering time and WUE chromosome 1 peak. 38
Supplementary Figure 20. Association of gene expression with key traits. 39
Supplementary Figure 21. Prediction of trait response to selection. 40
Supplementary Figure 21. Gene editing of diverse Arabidopsis accessions. 41
Supplementary Figure 22. FLC sequence coverage across accessions. 42
Supplementary Figure 23. Relative expression of FLC across accessions. 43
Supplementary Figure 24. Comparison of relative expression across exons. 45
Supplementary Figure 25. Summary of δ13C data from mutants and wildtypes. 46
Supplementary Figure 26. Correlation of mutant flc δ13C [‰] values and those from 19. 47
**XI .References** **48**

#### Supplemental Text: Extended text on analyses and results

##### I. Trait data

##### I.1 Data curation

The curated database of 1,862 traits comes from a combination of 108 laboratory and field experiments that use a subset of the 1001 Genomes *A. thaliana* accessions [1]. 228 traits come from the Arapheno database [2], and the remainder were compiled by Y. Voichek, D. Weigel, and M. Exposito-Alonso from published literature (**Table S1**). The trait data are averages for a genotype across replicates measured in replicated environments and were not measured in a single experiment or for single plants. Traits for different fitness data from various experiments are included, but for clarity, the primary fitness data used in the analyses comes from outdoor field experiments in two locations, Spain and Germany, with two rainfall and planting density treatments [3]. We classified every trait into a general functional category, and as being related to one of three drought adaptation strategies; *escape* (n=125), *avoidance* (n=402), and *tolerance* (n=756) [4] (**Fig. S1A, B**). We classified *escape* traits as those that can contribute to germination, growth, growth rate, vernalization, flowering time, and reproduction such as dormancy and germination. *Avoidance* traits were classified as those that can both help to conserve and increase access to water, including root growth and angle related traits, leaf area, biomass accumulation, stomatal density and size, and $\delta C^{13}$ (commonly used as a proxy for WUE). *Tolerance* traits were annotated as those involved with enduring low water content such as increased production of osmolytes and other metabolite related phenotypes, as they may play an important role in desiccation tolerance and osmotic regulation. While *tolerance* may be an important ecological strategy for *A. thaliana* in another context, we do not focus on these traits in this study as a bulk of them are metabolite traits or not life-history traits involved in ecological adaptation, and if they are involved in adaptation, it is not well understood. More focus in the future could be focused on these tolerance traits and metabolomics.

##### I.2 Water Use Efficiency

WUE is generally defined as the efficiency in which $CO_2$ is fixed relative to water loss and is instantaneously calculated as the ratio of net $CO_2$ assimilation ($A_n$) to transpiration ($E$) rates, two processes highly regulated by stomatal conductance [5]. Measuring instantaneous WUE is difficult in a single experiment because of the time required to measure gas exchange rates accurately for hundreds of individuals. However, overall or lifetime WUE can be characterized using a proxy measure $\delta C^{13}$, the ratio of carbon isotope $C^{13}$ to $C^{12}$ found in the leaf relative to atmospheric levels, as it is more fit for high throughput phenotyping. When stomata are closed, $C^{13}O_2$ is more likely to be assimilated by RuBisCo than when stomata are open, and thus the more often stomata are closed the leaf's isotopic composition will have an overall higher ratio of $C^{13}/C^{12}$ leading to higher $\delta C^{13}$ and water use efficiency. This ratio captures the leaf's history of closing stomata to avoid water loss and represents a mechanism of dehydration avoidance. Empirical studies have confirmed that $\delta C^{13}$ is correlated with (A/E)[5–7], assuming similar mesophyll conductance [8] and boundary layer conductance [9] (hereafter WUE refers to measurements of $\delta C^{13}$).

##### I.3 Trait imputation

The R package *missForest* was used to fully impute the trait dataset [10], as the traits had typically been only measured in a subset of accessions, with different subsets for different traits (**Fig. S1C**). This was done iteratively for each trait and all 1,135 accessions of the 1001 Genomes resource, with all other traits being used as predictor variables for each trait as a response variable. The algorithm was run with the default parameters, as attempts to increase the number of trees in the forest, or the number of variables considered at internodes did not improve the accuracy. We estimated the out-of-bag prediction error for each trait as the mean square error for predictions made in the out-of-bag sample that was 1 ⁄ 3 the size of the total dataset. This resulted in an average normalized root mean squared error (NRMSE) of 0.195 (see **Table S2** for estimated NRMSE for all traits; 95% highest posterior density (HPD) (0.071, 0.36); **Fig. S1D**). A NRMSE of 0.195 indicates that the errors of the model predictions are 19.5% of the variability in the response variable, *i.e.* the trait being imputed, thus the lower the value indicates the better the prediction accuracy. The imputation was necessary for many of the subsequent analyses performed with the traits, such as the principal components analysis (PCA), estimation of selection coefficients, and multivariate GWAS. While useful for many multivariate analyses (e.g. with many traits, there is not enough overlap in the raw data to conduct some analyses), we advise users to employ this dataset for analyses with care, as it is no substitute for new measurements. Traits with very few observations will likely have imputed data points closest to the mean of the population. We also provide the dataset in its raw. unimputed format, and advise that trait correlations are analyzed both in the imputed and the raw (with pairwise missing data removal) when possible.

#### II. Trait analyses

##### II.1 Principal component analysis

Principal component analysis (PCA) was used to decompose the variation of traits in a subset of 515 accessions for which field fitness data was available from a hot and dry environment [3]. For this, specific traits related to seed dormancy, vernalization, germination, flowering time, leaf traits, roots, stomata, growth rates, and stress response were isolated (n=205). We then removed highly correlated traits (Pearson's $R^2$ > 0.7), resulting in a total of 64 traits (**Fig. S1F**). Traits were scaled as *mean* and *sd centered*, then used in the primary PCA (**Fig. 1A**). To ensure the pattern we observed with a subset of the accessions was consistent with the entire species, we also performed a PCA using all native Eurasion accessions from the 1001 Genomes project (n=999) (**Fig. S1E**).

##### II.2 PC climate models

Using *randomForest*, we constructed two climate models to predict the phenotypic landscape of PC 1 and PC 2 across the native Eurasian range of *A. thaliana* (n=999). To build the models, we separately used climatic data (24 monthly min and max temperature, 12 monthly precipitation, and 19 bioclimatic variables; 55 total) associated with the localities of the focal 515 accessions as predictors for response variables PC 1 and PC 2. These models used 1,000 decision trees each, considering 18 variables at each split in the trees. The out-of-bag prediction errors recalculated as NRMSE for the models predicting PC 1 and PC 2 were 0.141 and 0.116, respectively. We also estimated the accuracy of the *randomForest* models by performing a cross-validation (CV) experiment using two thirds of the data for training and one third for testing. We then estimated the accuracy of the models by correlating the predicted values with the true values for the respective PCs. This was done iteratively (n=100) to establish a distribution of CV prediction accuracy estimates which was 0.72 (0.65-0.78 95%HPD) when using climate to predict PC 1 and 0.56 (0.48-0.62 95%HPD) when predicting PC 2. Using these models, PC 1 and PC 2 values were predicted for every map grid locality in the Eurasian native range of *A. thaliana* from longitude 15°W to 90°E, and latitude 34°N and 65°N **(Fig. 1D)**, thus generally mapping how phenotypes were geographically distributed and the localities were visualized after normalization with the same hues as in **Fig. 1A.**

##### II.3 Trait associations

We measured how many of the 1,862 traits were correlated with 4 flowering time datasets, water-use efficiency (WUE), and growth rate using Pearson's correlation coefficient with a significance threshold of 0.05 and the non-imputed dataset (**Table S3**). For all trait combinations, we correlated all raw overlapping pairwise trait values for 1,135 accessions [1], but for visualization purposes we sometimes subset the data to the 515 accessions for which we had complete fitness data (**Fig. 1B**). We also assessed how some trait associations changed when considering confounding factors. Specifically for flowering time and WUE, in addition to the primary association (**Fig. 1B**), we also constructed a multiple linear regression model using flowering time and additional fixed covariates such as latitude and seven genomics principle components (PCs) to predict WUE. In this model, flowering time is still the most significant predictor of WUE ($p < 4\times10^{-5}$), while latitude is also a significant predictor ($p < 0.03$).

We downloaded the averaged historical (1970-2000) bioclimatic, temperature, precipitation, and predicted evapotranspiration rate estimates from WORLDCLIM 2.0 for all the localities associated with the 1001 Genomes accessions. We estimated Pearson's correlation coefficient between climate data and target traits (**Table S4**). We fit linear regression models of the climate data as a function of these target traits. We did this first exclusively with the climate data and traits, and then with genomic principal components (PCs) from the 1001 Genomes project accessions and latitude as covariates in the model. Likewise, we calculated composite variables such as mean maximum and minimum monthly temperature, mean monthly precipitation, and mean predicted evapotranspiration rate estimates. The relationship with mean maximum temperature (Pearson's $r_{flowering10\text{-}mean.max.temp.} = -0.15$, p-value < $5x10^{-4}$; **Fig. S2A**). Contradictory, high WUE of *A. thaliana* accessions is correlated with low annual precipitation in their locations of origin during the growing season (**Fig. S2B**). Correlation between mean precipitation January-July in Eurasian range localities and flowering time at 16°C (Pearson's $r^2 = -0.14$, p-value = 0.002). In a model of flowering time at 16°C ~ mean precipitation + latitude + 4 genetic PCs, the coefficient for mean precipitation was estimated to be -0.122, p-value = 0.081 (**Fig. S2C**).

##### II.4 Climate change projections

We downloaded the averaged historical (1970-2000) bioclimatic variables [11] for the Eurasian range of *A. thaliana* at 5 $km^2$ (2.5 minutes) resolution. Predictions of future climate were retrieved from the Max Planck CMPI5 IPCC model projections of a business-as-usual scenario [11]. We plotted the 1970-2000 average value of annual precipitation and mean annual temperature as the beginning of the arrows (**Fig. 2E**) and the 2050 predicted values as the end of the arrows to indicate the trajectory of the climate in precipitation and temperature space. Arrows indicate whether the Euclidean distance (of both temperature and precipitation) between the current and 2050 climates becomes more similar to the current Spanish climate (red) or less similar (blue) (**Fig. 2E**). Trajectories with predicted precipitation and temperature more extreme than the Spanish site are in gray. The intensity of the trajectory is then mapped back to the initial localities across the Eurasian range to visualize where in the native range climate trajectories become closer to a Spanish climate or away from it (**Fig. 2F**). Note that while this map is a general climate prediction for whether the climate will be more or less similar to hot and dry Spain, it is not a comprehensive prediction of how climate will change across Eurasia.

#### III. Selection estimation

##### III.1 Total selection ($s$)

We computed total selection coefficients $s$ [12] using the covariation between relative lifetime fitness $w$ and trait variation $z$ (mean and variance centered): $w = sz$, where $s$ is the total selection coefficient. Total selection was first calculated on all non-fitness traits (n=1,823), as well as the first 20 PC axes, as they explain 75% of the phenotypic variation. For each trait, we computed $s$ with 100 bootstrap replicates with replacement from the trait data. Traits with a significant $s$ were determined as those with estimates whose 95%, 99%, and 99.9% CI does not overlap with 0, denoted as *, **, and ***, respectively. In order to compare these measures of selection, all fitness and trait data were mean and variance centered prior to calculating $s$. While there is fitness data from several studies, the largest collection of fitness data across accessions and thus primarily used in our selection analyses comes from an outdoor common garden experiment in two different environments, Madrid, Spain (m) and Tübingen, Germany (t) (**Figure 1E**), both with two rainfall treatments at each location, high (h) and low (l), as well as two planting densities, high/population (p) and low/individual (i) [3]. This resulted in a combination of eight different treatments denoted in this notation: mhp, mhi, mlp, mli, thp, thi, tlp, tli. For each treatment, three fitness traits were measured: survival to reproduction (as proportion of surviving replicates), seeds set (as the total number of estimated seeds set per plant), and lifetime fitness (as the number of offspring including those with suffered mortality and thus have 0 offspring). All fitness data were used to estimate total selection coefficients for all traits (**Table S5**).

The correlation of PC1 and fitness was clear in the most stressful hot-dry-high-plant-density environmental condition in Spain ($s_{fitness}$ = -0.152, $p_{boot} < 1.0 \times 10^{-3}$; **Table S6**), and was also clear in individual correlations of the major traits underlying PC1. In the flowering time data from the same study as the fitness data [3], we consistently observed that flowering time was under negative total selection for nearly every measure of both lifetime fitness and survival. If not significantly negative, then we found no association but never a positive one (**Table S7**). We note an exception to this trend in flowering time, though. The flowering time data from other studies also consistently shows flowering time under negative total selection, where there is only evidence of significant negative total selection on flowering time, or no association with fitness and flowering time (**Table S8**). Growth rate data showed selection consistently for fast growth ($s_{fitness}$ = -0.176, $p_{boot} < 1.0 \times 10^{-3}$; see all estimates in **Table S9**). There is consistent selection for high seed dormancy ($s_{fitness}$ = 0.129, $p_{boot} < 5.0 \times 10^{-2}$; see all estimates in **Table S10**). Finally, there is evidence of negative selection on WUE ($s_{survival}$ = -0.091, $p_{boot} < 5 \times 10^{-2}$; see all estimates in **Table S11**). This corroborates some of the aforementioned published results from outdoor field experiments [13–15] that suggest there is generally selection for early flowering time, the *escape* strategy, and as we show the traits that are also correlated with PC 1 and more generally, flowering time.

In-depth analysis of PC1's associations with all fitness measures in the hot-dry environment revealed the striking switch in correlation from negative when utilizing only survival as fitness ($s_{survival}$ =

-0.239, $p_{boot} < 1x10^{-3}$), to positive utilizing only fecundity as fitness ($s_{fecundity} = 0.109$, $p_{boot} < 5x10^{-2}$, **Table S6**). We also observe this pattern with flowering time from the original study where survival suggests selection for early flowering (**Fig. 1F**), but fecundity suggests selection for later flowering (**Fig. 1G**). This observation coincides with the total selection estimates for flowering time, where the estimates using fitness and survival show negative selection in the hot and dry environment, but estimates from seeds set show that flowering time is under positive selection ($s_{fecundity} = 0.085$, $p_{boot} < 5x10^{-2}$, **Table S7**). This pattern was also apparent in the trait growth rate, which was under significant negative selection with survival and fitness indicating selection for low absolute growth (escapers), but the estimates became nonsignificant when using fecundity (**Table S9**). Likewise, there was significant positive selection for dormancy when using survival and fitness data, indicating selection for high dormancy (escapers), or those that remain dormant as seeds throughout winter and germinate in early spring. When using fecundity data however, selection was significantly negative on dormancy (**Table S10**).

This switch in the sign of the total selection coefficients, in PC axis 1, flowering time, growth rate, and dormancy are all evidence insinuating the potential conflict known in ecology where later phenology increases the ability of plants to accumulate resources for reproduction but risk an increased mortality [16]. It could also be evidence that there are indirect trait effects impacting the strength and direction of selection in the hot and dry environment [17].

**III.2 Direct selection (β)**

To understand the direct effects of natural selection on traits, we accounted for correlations amongst traits using the multivariate selection approach: $\beta = P^{-1}s$, where $P$ is the phenotypic variance-covariance matrix [12], $s$ is the previously quantified total selection coefficient (without accounting for correlations), and β is the gradient of selection (or direct selection) that captures the independent effects of each trait that cannot be attributed to their associations with other traits. Through a series of trait decorrelation, and expert selection, we narrowed the analyses to 12 focal traits related to both seasonal and drought adaptation (with a minimum coverage of 43 accessions, and mean of 180) (**Table S12**): ABA accumulation in leaf (ref. [18]), WUE measured as δC13 (ref. [19]), primary dormancy (DSDS50; ref. [20]), secondary dormancy based on germination after 8 days in ℃ (d8_10C_perc; ref. [21]), flowering time at 16 ℃ (ref. [1]), absolute growth rate and relative growth rate (RGR) (ref. [22]), flowering after long day and vernalization (ref. [23]), root horizontal index and root RGR (ref. [24]), and stomatal density and size in fully developed leaves (ref. [19]) (see trait correlation matrix **Fig. S3**). We used all fitness data from ref. [52] (**Fig. S4**), but later focused the results to the hot and dry environment (ml) (**Table S13)** and the cool and wet environment (th) (**Fig. S5, Table S14)**. As before, we performed 100 bootstrap replicates with replacement to establish significance of β (direct selection) estimates.

When studying the consistency between total and direct selection over traits in the hot and dry environment (**Table S13**), we identified a primary congruent negative direct selection pressure over growth rate ($s_{fitness} = -0.185$, $p_{boot} < 1.0x10^{-3}$; $\beta_{fitness} = -0.209$, $p_{boot} < 1x10^{-3}$), where selection favors low absolute growth rates, suggesting selection for the *escape* strategy. However, in addition we found an incongruent direct selection over late flowering time (*avoider*s*)* once we accounted for trait correlations ( $\beta_{fitness} = 0.184$, $p_{boot} < 1x10^{-3}$; **Table S13**), even with survival ($\beta_{survival} = 0.19$, $p_{boot} < 1x10^{-3}$) (**Fig. 1H**). While a positive $\beta_{fitness}$ for flowering time may be due to the influence of fecundity on lifetime fitness, the positive $\beta_{survival}$ reveals there is also a survival advantage in addition to larger seed sets. These analyses are evidence that the apparent correlation of fitness with early flowering is a spurious one that not only vanishes but reverses when accounting for trait correlations.

We also observe selection for the lack of a vernalization requirement (**Fig. 1H**), more specifically negative selection on growth after vernalization, which marks additional evidence of selection for the *escape* strategy. Finally, in the hot and dry conditions, there was consistent positive total and direct selection estimated for root relative growth rate, or how fast roots grow relative to their current mass. While this trait axis lies orthogonal to our focal axis of trait variation (Fig. 1A), it suggests there may be benefits to this phenotype in dry conditions and this phenotype is predicted to be throughout Southern Eurasia.

In order to understand the influence of the indirect effects on flowering time, as well as the other traits, we performed a leave-one-out estimation to see if the removal of a single trait influenced the direct effect of selection on flowering time. After removing a single trait and performing the analyses an additional 12 times, showed that no single trait removal removed the positive direct selective effect on flowering time, however we did find that only when removing both absolute growth rate and growth after vernalization, did the significant positive direct effect on flowering time cease to exist (**Fig. S6**).

WUE, which counterintuitively strongly correlated with low survival in a hot-dry environment at both low plant density ($s_{survival}$ = -0.112, p-value$_{boot}$ < $1x10^{-3}$) became non-significant while accounting for trait correlations (**Table S13**). Moreover, WUE in Germany, which was non-significantly related with survival in well-watered conditions (**Fig. S5, Table S14**), became positive in the experimental treatment in Germany where a Spanish low precipitation was imposed ($s_{survival}$ = 0.027, p-value$_{boot}$ < $5x10^{-2}$; even after correcting for flowering time effects, $\beta_{survival}$ = 0.075, p-value$_{boot}$ < $1x10^{-3}$; **Fig. S7**). These analyses provide evidence of a conflict in natural selection between two adaptive strategies to water limitation in realistic outdoor climates that may lead to inefficient phenotypic evolutionary responses [16,25].

We also used fitness data from additional common garden experiments in varying environments [26,27] to assess what patterns of selection have been observed in other common gardens for these same traits. In the Fournier-Level et al. (2011) data, we see that in the Finland environment there is selection for high absolute growth rate and slow relative growth rate (RGR), suggesting selection for *avoiders.* Likewise, in Germany slow absolute growth and fast RGR were selected for, suggesting selection for *escapers*. Flowering time does not appear to be under strong direct or total selection. Interestingly, in the Finland environment, WUE is under negative selection (**Fig. S9**). Manzano-Peidras *et al.* 2014 found evidence for selection for late flowering, but our analysis shows that when correcting for correlated traits, there is a negative direct selective effect on flowering time (**Fig. S10**), indicating that in this experiment and in these populations there may also have been a selection conflict on flowering time.

**IV. Genome wide associations**

All trait data with continuous values were normalized via quantile transformation as using quantile-transformed trait data in GWA has been shown to reduce type 1 error rates compared to non-normal, untransformed values [28]. For count data and traits with many zero values, we first performed a log(x+1) transformation, followed by a quantile transformation. Binary phenotypes were not transformed. We assessed how the correlation structure of the trait data was impacted by both the imputation (I.3 Trait imputation) and quantile normalization (**Fig. S11**).

**IV.1 Linear mixed models**

To perform genome wide associations (GWA), we started with the 1,135 accessions and 11,769,920 SNPs from the 1001 Genomes Project. Depending on trait data coverage, not all accessions and SNPs were used in every GWA. We ran subsequent GWA on first the 12 focal traits, but then also all traits using the raw data, the normalized data, and finally the imputed data. For each trait we conducted GWA using a univariate linear mixed model (LMM) implemented in GEMMA (ref. [29]). This model corrects for genome-wide background differentiation collinear to the phenotype of study (i.e. relatedness and population structure correction) with a kinship matrix. IfIf minor allele frequency (MAF) (MAF) or missing data cutoff werewere used, this was explicitly stated. Measures of SNP-based heritability for each trait were estimated using the random effect of LMM informed by the kinship matrix, which quantifies how much genome-wide relatedness explains the variation in the phenotype.

Following GWA with the various trait datasets, we compared the estimates of effect sizes across the datasets using Pearson's correlation coefficients to assess how the data imputation and normalization impacted the ranks of effect size estimates (**Fig. S12**), but it did not appear to drastically impact the estimates, though we know there was an impact, the biggest being between raw or normalized data with imputed data, as we know the imputation changes the underlying trait variance (**Fig. S11**).

SNP-based narrow sense heritability estimates are estimated in GEMMA as the total trait variance explained by additive genetic variation [29]. These estimates are sensitive to the number of individuals used, as the estimate is based on the total trait variation and genetic variation which will always change slightly when different accessions are used, in this case, each GWA was run with a subset of the inbred 1001 Genomes Project accessions. Heritability was estimated for all traits in the BSLMM GWA (see below), and the LMM GWA for both normalized and raw trait data. When calculating mean heritability, we did not consider estimates that had a 95% HPD interval that spanned over 80% of the distribution of heritability, and in the case of LMM, we did not include estimates that had a standard error greater than 0.25. Overall we found that the most traits had moderate-to-high SNP-based heritability $h^2_{BSLMM}$ = 0.55 [0.25-0.85; n = 1,275], $h^2_{norm}$ = 0.34 [0-0.72, n = 1,251], $h^2_{raw}$ = 0.38 [0-0.80, n = 1,255]. Additional parameters from BSLMM indicated many traits were polygenic as well with an average of $\gamma n$ = 48.5 causal loci [21.6-75.4] (**Table S15**). We also found that *escape* traits had higher heritability estimates (mean $h^2$ = 0.71 [0.43-0.99]) than *avoidance* traits (mean $h^2$ = 0.49 [0.03-0.94]) (wilcox test p-value=$2.6\times10^{-8}$, **Fig. S13**).

**IV.2 Bayesian sparse linear mixed model**

We subsetted the original imputed SNP matrix from http://arapheno.1001genomes.org with 10,709,466 SNPs to 1,353,386 SNPs for all 1,135 accessions to be used in a Bayesian sparse linear mixed model (BSLMM) GWA, implemented in the software GEMMA (Zhou and Stephens 2014), for each of the 1,862 phenotypes. This model estimates more parameters from the data and therefore takes a long time with many SNPs, therefore subsetting to higher quality SNPs across the genome. The BSLMM full model is a derivation of LMM, but unlike LMM, BSLMM implements two random effect distributions: effect sizes are either attributed to direct effects, that is, those that directly affect the trait and are normally distributed with broad variance, and indirect effects, that is, background effects that all SNPs have with a narrow variance. In this way, the number of major effect loci, $\gamma n$, are estimated along with the trait variation explained by the major effect loci (PGE), as well as all additive genetic variation (PVE, or narrow-sense heritability). Additionally, a hyperparameter, $\rho$, is estimated to approximate PGE, except here a low value of $\rho$ indicates the genetic architecture is fit for a LMM as the trait is polygenic, however a high value of $\rho$ close to 1 indicates the trait has few large effect loci, more fit for a Bayesian variable selection regression (BVSR) model. BSLMM is a hybrid model that can accommodate both types of genetic architectures and estimate parameters summarizing genetic architectures across traits (**Table S15**).

**IV.3 Multivariate linear mixed models**

For the 12 key traits used in the selection analysis (**Table S16**), pairwise multivariate LMM (mvLMM) GWA were performed [30]. Again, analyses were run with non-imputed and imputed data, and when the data were not imputed only 16/66 pairs of mvLMM GWA were possible given lack of overlapping samples and large number of parameters to estimate. These were run with an allele frequency cutoff of 0.05, 5% missing genotype data maximum, and correlated effects of SNPs were corrected for using a kinship matrix. In the case of flowering time and WUE, there were 248 overlapping samples with trait data for both traits, and after passing thresholds, 546,668 biallelic SNPs remained and were associated with the two traits via a mvLMM (**Fig. 3A**). We also re-estimated these effect sizes using an additional five genetic PCs as covariates in the model to control for additional background population structure in the model (**Fig. S15)**; this marginally improved QQ plots (**Fig. S16**). For flowering time and growth rate, there were 395 overlapping samples, and 347,257 SNPs were used (**Fig. S17**). As mentioned, in addition to the 16 pairs of traits with enough accession overlap, we also conducted mvLMM GWA for all 66 pairwise trait combinations using the imputed trait data. We compared the genetic correlations estimated from the approaches using imputed and non-imputed data (**Fig. S18**), and for the comparable pairs the estimates were not drastically different, in fact the genetic correlation estimated between flowering time and WUE decreased when estimated with imputed data.

To further investigate the phenotypic effects on flowering time and WUE of specific genomic loci we investigated a highly significant novel locus in chromosome 1 that had not been described before (this locus was consistently a top hit in GWA with imputed [n=1,135 accessions] and non-imputed trait data [n=248 accessions], with pruning low frequency or poorly genotyped SNPs, utilizing kinship correction, and 5 genomic PC covariates, **Fig. S15**). The top SNPs in chromosome 1 span ~7.5 kb and overlap with five candidate genes (AT1G11500-AT1G11540) (**Fig. S19)**. While several are not fully characterized, AT1G11500 appears to be involved in leaf development and determination of bilateral symmetry, AT1G11510 encodes a transcription regulator, AT1G11530 encodes a protein with disulfide isomerase activity, AT1G11540 encodes a sulfite exporter TauE/SafE family protein involved in protein ubiquitination and is highly expressed in the root, and AT1G11520 has no known molecular functions (**Tables S17, S18**). Notably, in the mv LMM GWA from imputed flowering time and WUE data, only AT1G11540 and two additional genes contained highly significant alleles near the chromosome 1 peak, one associated with embryo development ending in seed dormancy (AT1G11680), and the other involved in a cellular macromolecule metabolic process (AT1G11740) (**Tables S19, S20**). The expression of three of these genes correlates with flowering time and expression of AT1G11540 correlates with WUE (**Fig. S20**) (transcriptome data from ref. [31]).

**IV.4 Genetic correlation approximations**

In addition to estimates from mvLMM GWA, we also approximated genetic correlations among traits using Pearson's correlation of summary statistics of SNPs from univariate GWA. We summarized effect sizes of SNPs as Z scores where $Z = (\frac{\beta}{se})^2$ . For this approximate genetic correlation, we only correlated unlinked SNP effects by randomly selecting SNPs from independent linkage blocks that had high confidence variant calling, resulting in ca. 60,000 SNPs. With these approximate genetic correlations, we assessed how consistent they were with genetic correlation estimates from mvLMM GWA (**Fig. S18**). It is hard to say whether the genetic approximation correlations are as accurate as those from the mvLMM, as few estimates were possible with raw data and the imputed estimates could be inflated, however the key genetic correlations between flowering time, growth rate, and WUE all appear consistent regardless of the estimation process (**Fig. S18**).

**IV. 5 Fitness effects**

Fitness data from outdoor experiments of 515 *A. thaliana* accessions [3] were used in a univariate LMM GWA with no MAF cutoff and corrected for population structure using a kinship matrix. The sign of the effect size estimates for each allele indicated whether the allele has a positive or negative inferred effect on the fitness measure. We use these effect size estimates to color alleles to indicate positive (green) and negative (red) selection (**Fig. 3A, B, C**). In an attempt to account for the variation in fitness that is explained by the variation in growth rate, we fit a linear model of fitness explained by growth rate. The residuals of this model represent the remaining variation in fitness not explained by growth rate, and we therefore used these residuals in an additional LMM GWA, with the same parameters as above, to assess the remaining selection pressure on the alleles after accounting for selection on growth rate (**Fig. 2A**).

For visualization purposes, the SNPs with top 0.05% effects on both flowering time and WUE (from **Fig. 2A**) were plotted against the expected allele frequency change ($\Delta p$) based on their correlation with lifetime fitness (**Fig. 2C**). We then used as $\Delta p = p(1 - p)a$, where $p$ is the starting allele frequency, and $a$ is the estimated allele effect in relative fitness based on the effect size estimates from the GWAS of fitness.

**V. Prediction of phenotypic response to selection**

We projected the population phenotypic responses to natural selection by using the Breeder's Equation or selection response equation (ref. [49]) $\Delta z = GP^{-1}s$, where $G$ is the additive genetic variance-covariance matrix constructed from heritability and genetic correlation estimates among the key 12 traits, $P$ is the phenotypic variance-covariance matrix, and $s$ is the raw covariance of the trait with lifetime fitness in hot-dry conditions. For this, we used the estimates of $s$ from lifetime fitness in both the hot-dry environment (Madrid (m) and low water (l)) and the cool-wet environment (Tübingen (t), high water (h)), for both individual (i) and population (p) planting densities. For the genetic variance-covariance matrix, we used the SNP-based heritability estimates from key traits (**Table S16**), and the genetic correlations estimated from multivariate GWA (**Fig. S18**). Estimates of phenotypic change are reported for the two environments (**Table S21, Fig. S21**).

#### VI. Gene editing of FLC

##### VI.1 Establishing lines

To test the effects of attenuating *FLC* activity in different accessions of *A. thaliana* with a wide range of functional *FLC* alleles, we selected 60 late- and two early-flowering accessions for CRISPR/Cas9 gene editing. The accessions were chosen from the 1001 Genomes Project [1]. Based on the published FT16 (16°C and long day photoperiod [16 h light / 8 h dark]) phenotypes (https://arapheno.1001genomes.org/phenotype/262/), late-flowering accessions were selected with an FT16 range of 60.3 to 104.0 days until first open flower. Two early-flowering accessions 7186 and 9908, producing the first open flower after 49.75 days and 46.5 days, were also included (**Fig. S21** and **Table S22**). The reference accession Col-0 (1001 Genomes id #6909) was included in most phenotyping experiments as a general comparator. The selected accessions cover a broad genetic and geographic range of the species, representing about 13 out of 25 major *FLC* haplotypes in the global population [1] (**Fig. S21B** and **Table S23**).

##### VI.2 CRISPR/Cas9 protocol

To generate *flc* mutants we chose a targeted forward genetics approach: Constructs expressing paris of gRNAs (0.1, 1.3, 12.3, 13.4) were transformed as pools into the 62 accessions, followed by a phenotypic screen for accelerated flowering in the T2 generation and counter-selection of the transgene to obtain a total of 112 lines (**Fig. S21C and D**). As both the gRNA target site and the genomic context are determinants for Cas9 efficiency [32,33], we selected these multiple target sites preferably not located in the *FLC* chromatin nucleation region to avoid potential Cas9 impairment deriving from methylation and chromatin compaction (induced during flowering-promoting and FLC-repressing vernalization) [34–39]. The gRNAs were designed using CCTop and no off-target binding was predicted [40]. Utilizing the 1001 Genomes variant data we made sure to select gRNA against conserved sites [1]. Only one accession (9583) had one SNP in the sequence of gRNA 13.4.

For CRISPR/Cas9 mutagenesis, we slightly modified a previously established toolset [41] with a supermodule destination binary vector carrying a plant-codon optimized Cas9 driven by a *UBQ10* promoter. According to [42], a TTTT stretch in the sgRNA scaffold reduces the efficiency of sgRNA transcription and consequently decreases the efficiency of genome editing. We aimed at mutating the TTTT stretch but also at introducing a new unique restriction site to facilitate the introduction of sgRNA target sites via oligo annealing. We found that mutating the TTTT stretch to TTCT introduced a new unique *XbaI* site, therefore achieving both goals. This unique *XbaI* restriction site was introduced into the D and E modules from ref. [41] at position 2,405 by mutation-PCR [43,44]. D and E modules were linearized by double digestion with *FaqI* and *XbaI*. Universal and target-site specific oligos (**Table S24**) were phosphorylated with T4 Polynucleotide Kinase (Thermo Scientific, USA): 1 µl oligo (100uM), 0.5 µl T4 PNK, 1 µl T4 PNK buffer A, 1 µl ATP (10 uM), 6.5µl nuclease-free water, incubation at 37˚C for 40 min, and 75°C for 10 min. 0.5 µl of each phosphorylated oligo (at a concentration of 10µM) was combined and annealing was allowed by heating the oligo mix to 95°C and subsequent cooling to room temperature. Successful annealing was verified by electrophoresis on a 2% agarose gel. This mix was diluted 1:50 with nuclease-free water and 0.5 µl was used as an insert for ligation to the prepared linearized D or E backbones with T4 DNA ligase (Thermo Scientific, USA): 5 ng D or E backbone, 0.5µl annealed fragment, 0.5 µl T4 ligase, 1 µl T4 buffer, 1µl ATP (10µM). The mix was incubated at room temperature for 10 min and 2 µl were used to transform the *E. coli* strain DH5α. The following pairs of target sites were combined in each one supermodule, with the procedure described in ref. [41]: 0.1 (D) and 13.4 (E), 1.3 (D) and 13.4 (E), 12.3 (D) and 0.1 (E). The three resulting supermodules were pooled in a 1:1:1.5 ratio and this mix was transformed into the *Agrobacterium tumefaciens* strain ASE [45].

##### VI.3 Plant transformation

Seeds of all wild-types were sown in four pots each of size 13 x 9 x 5 cm. Seeds were allowed to germinate at 22˚C under long days (LD, 16 h light / 8 h dark) and after full expansion of the first true leaves, plants were thinned to keep eight to twelve plants per pot. Plants were vernalized for 12 weeks at 4°C and short days (SD, 8 h light / 16 h dark) to ensure saturated vernalization and abundant development of floral organs after the subsequent transfer to a greenhouse with cultivation at 22˚C under LD. At full development of inflorescences, accessions were transformed with the floral dip method [46] carried out twice with a 7 to 10-day interval. At full seed maturation, all $T_1$ seeds from all pots per accession were pooled.

Seeds from plants that were carrying the AT2S3::mCherry [41] cassette as T-DNA marker were screened for fluorescence or absence thereof under a LEICA MZFLIII Fluorescence stereoscope (Wetzlar, Germany) with a SOLA 365 SM Light Engine© lamp (Lumencor, USA). Up to 50 positive seeds were selected per accession. Up to 15 positive primary transformant seeds ($T_1$) were sown in one pot, with different numbers of pots per accession depending on the number of seeds selected. After plants were established at 22°C under LD, they were treated with a heat cycle to increase Cas9 editing efficiency according to [47]. After a five days recovery phase at 22°C under LD, pots were transferred to 4°C under SD for four weeks, for a non-saturating vernalization treatment to ensure the repression of residual *FLC* expression deriving from heterozygous or chimeric *FLC* editing in the primary transformants but to prevent full repression of *FLC* expression in plants with native, unedited *FLC*. The pots were then transferred back to 22°C under LD and the plants were allowed to propagate. Strong differences in flowering time were detected between the vernalized $T_1$ individuals, indicating that the vernalization treatment was indeed not saturating and sufficient to compensate for allelic heterogeneity. Seeds from all $T_1$ plants that reached the generative phase were bulked per accession to generate $T_2$ seed pools ($T_2$-pools). Note that the relative proportion of $T_1$ plants that did not flower at all differed between accessions.

#### VII. Characterize *FLC* mutants

##### VII.1 Mutant screening

We expected that successful editing of *FLC* and the inheritance of mutations would lead to early flowering in the $T_2$ generation across all accessions. To identify rare early flowering individuals in a large late flowering population, we sowed over 1000 seeds per 40 x 30 cm tray, assuming editing events to be infrequent. Early flowering individuals were visually screened after inflorescence development and transferred to individual pots for optimal growth and seed set. Approximately 25 early flowering plants per accession were propagated to the $T_3$ generation (totaling more than 2,500 plants from more than 120,000 screened individuals). $T_3$ seeds were examined for presence of the transgene using the fluorescence reporter system. Seeds from fully non-transgenic lines, or selected non-transgenic seeds from segregating lines, were chosen for further analysis. Flowering time was analyzed for one to six $T_3$ lines (five plants each) at a constant temperature of 22°C under LD conditions, alongside five wild-type plants. One plant from one to two lines per accession of early flowering, non-segregating lines was selected for sequencing.

The genomic DNA of *flc* KO or KD lines were sequenced using Illumina 2x150 bp reads. Raw reads underwent trimming with cutadapt, followed by mapping to the TAIR10 reference genome of *A. thaliana* using bwa-mem (version 0.7.17-r1188) [48]. Duplicate reads were eliminated with picard (version 2.18.25) MarkDuplicates. SNP and small insertion-deletion (Indel) calling were conducted with GATK (version 4.1.0.0) HaplotypeCaller, applying recommended best practices with some modifications [49]. Variant filtering employed GATK (version 4.1.0.0) VariantFiltration with the criteria "QD < 22.0 || FS > 10.0 || MQ < 50.0 || MQRankSum < -5.0 || ReadPosRankSum < -5.0". Variant effects were annotated using SnpEff (version 4.1l). To verify the correct identity of mutant backgrounds and wildtypes, SNPmatch (version 5.0.1) was employed [50].

**VII.2 Characterization of transgene insertions**

T-DNAs can insert incompletely or as chimeras due to border instability, leading to false positive and false negative transgene reporter signals [51–53]. Whole-genome sequencing showed that ten early-flowering lines (9%) indeed carried foreign DNA derived from the transgenes (**Fig. S22** and **Table S25**). To detect potential partial transgenic sequences not detectable by seed fluorescence, we aligned all Illumina short reads to the full-length 10,275 kb expression plasmid [41] using bwa-mem (version 0.7.17-r1188). Mapped reads were retained from the resulting bam files, and high mapping quality (MQ) was ensured using samtools (version 1.10) with "view -b -F 4 -q 30". Reads with mismatches were filtered out with bamtools (version 2.5.1) using "filter -tag NM:0". To identify coverage segments in the non-*A. thaliana* (TAIR10) portion of the plasmid, per-base coverage information was obtained with samtools and "depth -aa". Sequences from *A. thaliana* (*UBQ10*, *AT2S3*, *U6*) in the plasmid served as controls for the presence of mappable reads. Clipped reads were extracted from these alignments using samclip (version 0.4.0) with "--invert --max 130". After converting resulting bam files to fastq format with bedtools "bamtofastq," the reads were remapped to TAIR10 using bwa mem. Only clipped reads were retained using samclip with "--invert --max 140." Regions with coverage in TAIR10 were identified with samtools and "depth." Insertion positions were determined from the region with the maximum coverage, treating sequences less than 150 bp apart as a single region. Positive controls were hits in regions of genes *UBQ10*, *AT2S3*, and *U6*, extending 2 kb upstream and downstream in all samples (Chr.3: 4559112-4563212; Chr.4: 2715977-2722308; Chr.4: 13609796-13614563). The insertion position in line ID5741-02-R03 could not be definitively identified due to the clipped reads mapping perfectly to multiple positions in the TAIR10 genome. Additionally, the positions of insertions in lines ID8283-01-R04 and ID8283-02-R04 were undetermined as the clipped reads (or subfractions) did not map to TAIR10, and a blastn search yielded no results. The line CS77069-08-R03 was initially misclassified as non-transgenic but was later confirmed to carry a single-copy, full-length transgene with confirmed seed fluorescence. We kept these lines in the sample set for experiments in contained environments but acknowledged the potential of second-site effects from the transgene.

CRISPR-induced SNPs and small indels at target sites 0.1 and 12.3 were extracted from annotated VCF files using vcftools in the chromosome 5 region spanning 3,173,700 to 3,173,900 bp for gRNA target site 12.3 and 3,179,600 to 3,179,750 bp for gRNA target site 0.1. Characterization of CRISPR-induced deletions larger than 25 bp at the *FLC* locus employed a sliding-window linear regression approach: Per-base coverage information was obtained using samtools, focusing on the region 3,162 to 3,188 kb of chromosome 5. Linear regression were then performed using per-base coverage against the genomic coordinate in a 25 bp sliding window and a step-size of 1 bp, with the window- and the step-size parameter determined by finding the best-performing ones from a series. Deletions were called as "flat valleys" of sliding-window mean sequencing coverage by scanning the slope value of the sliding window linear model. To avoid spurious calls, the mean sequencing coverage for the called windows were set to zero. The precise deletion boundaries were then identified by tracing back within the focal sliding windows to the first and last positions without sequencing coverage according tot per-base coverage data. Calls in the region 3,174.63-3,174.73 kb displaying natural genetic structural variation were excluded. Separation of deletion calls in some samples due to spurious mapping of single reads was addressed by transforming all per-base coverage values <= 2 to 0, concatenating separated deletion calls, and correcting start and end coordinates. The transformed data were used for Figure 3A. Variant analysis of a 9 kb genomic region spanning *FLC* uncovered a large variety of induced *FLC* mutations, mostly due to Cas9 activity at target sites 0.1 and 12.3 (Fig. 3A). 59 lines representing 40 accessions carried deletions larger than 25 bp with the largest deletion being 6,797 bp. 53 lines representing 37 accessions carried deletions smaller than 25 bp or SNPs, mostly deriving from editing activity at the 12.3 site, with mutations at the last intron-exon junction (Fig. 3A and **Table S26**).

##### VII.3 qRT expression analysis

Seeds were surface sterilized with 70% EtOH and plated on sterile medium with ½ MS (incl vitamins and MES, pH5.8, 1% agar-agar. Three biological replicates were distributed over three plates, each representing the wildtype and one or two mutants. The position on the plate was randomized. To reduce the effects of photoperiodic differences throughout the day and the circadian clock, photoperiods of three plant growth incubators (one for each set of biological replicates) were set at a shifted schedule of two hours. As the sampling of all samples of one replicate block took not more than 40 minutes, the total covered photoperiodic timespan of the experiment was also 40 minutes. The chambers were set at 22°C LD, and the sampling time point was ZT14 (two hours before dark). Several seedlings per line were pooled as one sample. RNA from each biological replicate was extracted individually using a column-based protocol adapted from [54]. RNA quality was validated with a Nanodrop spectrophotometer and normalized to 500 ng/ul. 1 µg total RNA was reverse transcribed with RevertAid First Strand cDNA Synthesis Kit (Thermo Fisher Scientific, Waltham, USA) using an oligo (dT) primer. The cDNA equivalent of 30 ng total RNA was used in a 15 µl PCR reaction with SYBR Green (Thermo Scientific Maxima SYBR Green qPCR Master Mix) in a CFX384 Real-Time System Cycler (BioRad, München, Germany). Run- and plate-specific effects were normalized using some control samples replicated on all plates. The relative quantification was calculated with the ΔΔCt method using *ACT8* (AT1G49240) as a standard [55] and calibrated by biological replicate 1 (rep1 ) of Col-0.

For *FLC*, some samples yielded no usable Ct values (Ct > 38). As the same samples showed consistent expression of *ACT8* and investigated other genes, we assumed that the expression of *FLC* in these samples was extremely low and therefore undetectable. The samples of Col-0, which does not express *FLC* due to a trans-regulatory defect in *FRI*, and lines with full length *FLC* deletions (having lost specific priming binding sites) served as *FLC*-null controls to determine the lower, false-positive noise signal of relative expression 5 in the qRT reaction that was likely generated through unspecific amplification during the very late stage of the qPCR reaction, possibly due to the high sequence homology between genes encoding FLC related MADS box transcription factors [56,5758–60] (**Fig. S23**). For line ID9599-02-R02, only the *FLC* transcript level was determined. Truncated *FLC* transcript was identified by fitting a linear model FLC_expression_exon_7 ~ FLC_expression_exon_2/3, extraction of model residuals and comparison of each mutant to the respective wildtype (**Fig. S24**). All primer sequences are listed in **Table S24**.

##### VII.4 Measurement of carbon isotope ratio

Mutants and wild types were grown in three pots representing three biological replicates at 22°C under LD. The pots were randomly distributed over a total of ten trays, which were rotated and moved every day to reduce position effects. After allowing germination and establishment of the first true leaves, plants were thinned to three plants per pot. The leaves of different plants never overlapped with each other during the experiment. Rosettes were harvested at the initiation of flowering or after 22 days after germination (whatever was first). Depending on the rosette size at the initiation of flowering several plants were combined to one replicate to reach the required amount of tissue for analysis. Plant tissue was dried at 60°C for 24 hours and homogenized in 5 ml tubes containing 5 ball bearings to very fine and uniform powder. Dried material was transferred to a 1.5 ml microfuge tube and sent to Isolab GmbH (Schweitenkirchen, Germany) for an analysis of carbon isotope composition ($\delta^{13}C$) with $^{13}C$-CF-IRMS. Four technical replicates per sample were analyzed. For a more detailed description of the procedure, see [61]. Data are presented as $\delta^{13}C$ [‰] vs. V-PDB (**Supplementary Fig. 25C**).

Utilizing this novel resource, we measured carbon isotope composition ($\delta^{13}C$ [‰]) as a proxy for WUE in a subset of 20 wild-type accessions (including Col-0) and 29 *flc* mutants (with a maximum relative *FLC* expression level of 18.3 a.u.), which included 19 *flc* mutant / wild type contrasts. As expected, the *flc* mutants flowered earlier (mean [se] = 22.1 days [1.0]) than the wild-type parents (mean [se] = 73.3 days [6.7]) and had lower $\delta^{13}C$ levels**:** In pairwise comparisons with the respective wild type, nine *flc* mutants representing five genetic groups showed significantly lower (more negative) $\delta^{13}C$ levels (two sided Student's t test, Benjamin-Hochberg correction, p.adj. < 0.05) (**Supplementary Fig. 25B)**. The difference in $\delta^{13}C$ ($\delta^{13}C_{diff}$ [‰]= $\delta^{13}C_{Wildtype}$ [‰] - $\delta^{13}C_{Mutant}$ [‰]) in these nine contrasts was between 0.55 ‰ and 1.28 ‰. It is important to note that while some samples showed high variance between biological replicates and therefore the reduction of $\delta^{13}C$ did not pass the significance test, the *flc* mutant derived from accession 9421 was clearly unaffected in $\delta^{13}C$ levels (**Supplementary Fig. 25B)**.

#### VIII. Growth chamber experiment with FLC mutants and wildtypes

50 *flc*-derived lines, along with 50 wild-type lines, were grown in two parallel experiments simulating a hot environment with no water limitation (16h light, ca. 1000 Lux, and 21℃) to quantify flowering time and fitness through seed yield. First, seeds were stratified at 4ºC in Eppendorf tubes with a small amount of water to break any dormancy on the seeds. Seeds were planted in trays with 24 pots (5.72cm x 7.26cm x 6.35cm each). Three seeds were planted per pot but were thinned to one plant per pot after one week in a standard soil mix. As there was no water limitation, trays were bottom watered approximately twice a week and occasionally more if the top of the soil appeared dry. Flowering time was measured as days to first petal emergence and seed weight was measured as total weight of seeds harvested (mg).

##### VIII.1 $\delta^{13}C$ imputation

In order to analyze $\delta^{13}C$ for the 50 wild-type accessions, we relied heavily on the $\delta^{13}C$ data that already exists for several hundred accessions [19], and then our imputed dataset of $\delta^{13}C$ data NRMSE = 0.13; **Table S2**) that filled in for any remained accessions without $\delta^{13}C$ data. For the *flc* mutants, we used the 19 values of $\delta^{13}C$ measured above, and then imputed the values for the remaining 31 accessions using those 19 values, the $\delta^{13}C$ data from [19] (see correlation **Fig S26**), and the flowering time of the accessions from this experiment in order to ground the values predicted to this experiment and not general values of $\delta^{13}C$. Likewise, for both the wild-type and mutant $\delta^{13}C$ data, we scaled the values using $\delta^{13}C$ values from Col-0 grown in similar conditions, thus making this a measure of WUE relative to Col-0.

#### IX. Supplementary Tables

**Supplementary Table 1.** Summary of trait data assembled in this study.
**Supplementary Table 2.** Trait imputation accuracy.
**Supplementary Table 3.** Trait associations with flowering time, WUE, and growth rate.
**Supplementary Table 4**. Significant associations between target traits and climate variables.
**Supplementary Table 5**. Total selection coefficients estimated for all non-fitness traits.
**Supplementary Table 6.** Total selection coefficients estimated for PC axes 1 and 2.
**Supplementary Table 7.** Total selection coefficients estimates for all flowering time data.
**Supplementary Table 8.** Total selection estimates for flowering time traits from other studies.
**Supplementary Table 9.** Significant total selection estimates for growth rate related traits.
**Supplementary Table 10.** Significant total selection estimates measured for dormancy traits.
**Supplementary Table 11.** Significant total selection estimates for WUE ($\delta C^{13}$).
**Supplementary Table 12.** Overlap of trait data with corresponding fitness data.
**Supplementary Table 13.** Total and direct selection estimates in the Madrid, low water site.
**Supplementary Table 14.** Total and direct selection estimates in the Germany, high water site.
**Supplementary Table 15.** Heritability and GWA parameter estimates for all traits.
**Supplementary Table 16.** Heritability and GWA parameter estimates for 12 key traits.
**Supplementary Table 17.** GO annotations for flowering time and WUE.
**Supplementary Table 18.** GO annotations for flowering time and WUE; with 5 genetic PCs.
**Supplementary Table 19.** GO annotations for flowering time and WUE; imputed data.
**Supplementary Table 20**. GO annotations for flowering time and WUE; imputed data with 5 genetic PCs
**Supplementary Table 21.** Selection response predictions.
**Supplementary Table 22.** Accession information for lines used in gene editing.
**Supplementary Table 23.**The 25 major FLC haplotypes.
**Supplementary Table 24.** Universal and target-site specific oligos for FLC**.**
**Supplementary Table 25.** Summary of foreign DNA derived from the transgenes in lines.
**Supplementary Table 26.** Annotation of FLC mutations across diverse accessions.
**Supplementary Table 27.** Transcript analysis of FLC.

#### X. Supplementary Figures

**Supplementary Figure 1.** Characterization of trait data.

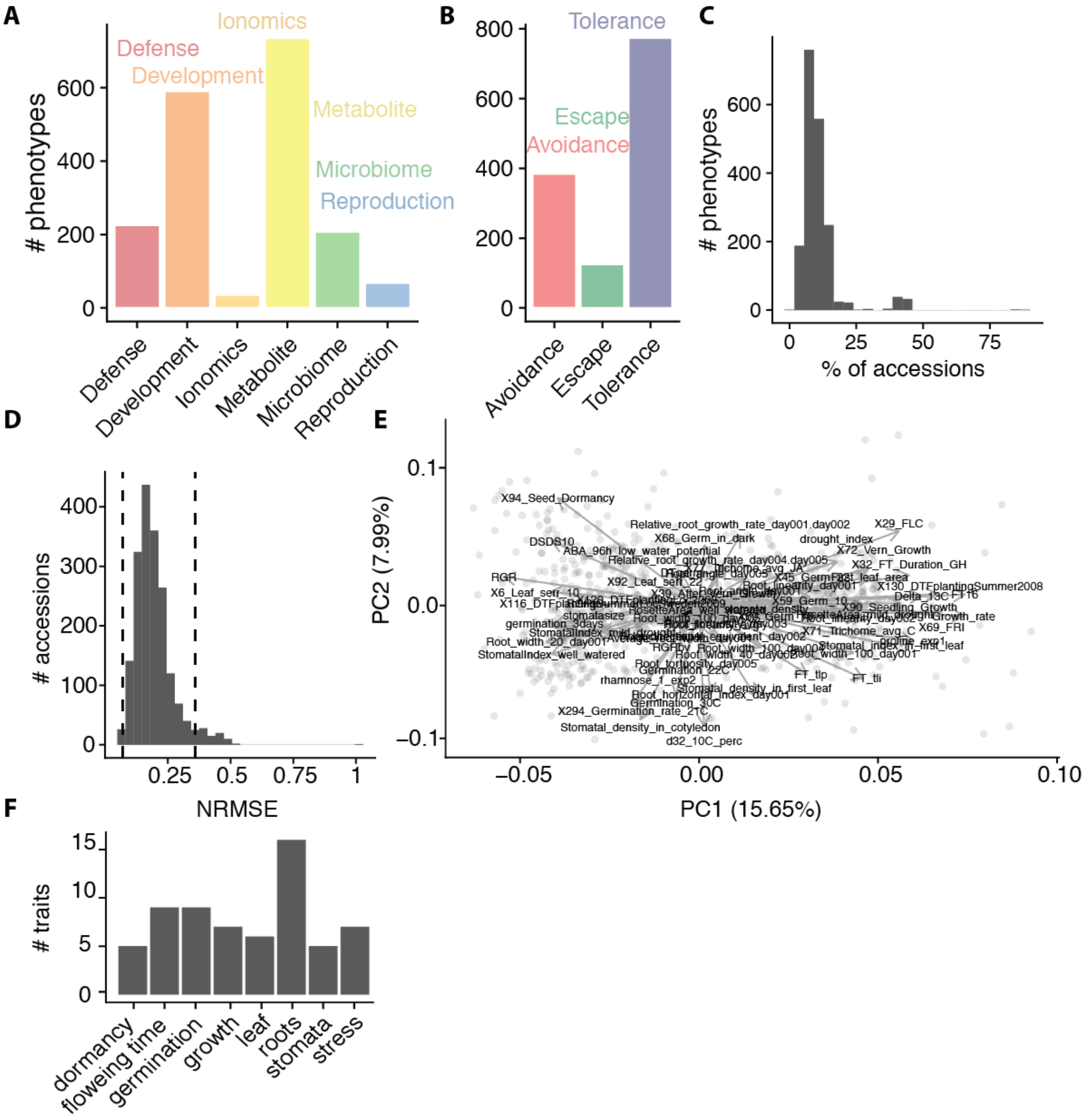


**(A)** Raw counts of traits classified into general functional categories and (**B**) drought response strategies. (**C**) Coverage of 1862 traits across the 1001 Genomes accessions (n=1,135). (**D**) Histogram of Normalized root-MSE for all traits (n=1,862) that were imputed. (**E**) PCA axes 1 (15.6%) and 2 (7.9%) of 64 target phenotypes for all Eurasian accessions of *A. thaliana* (n=999); trait associations are in the same direction as in Figure 1A. (**F**) Counts of 64 traits in different functional trait categories used in the principle components analysis.

**Supplementary Figure 2.** Climate associations for flowering time and WUE.

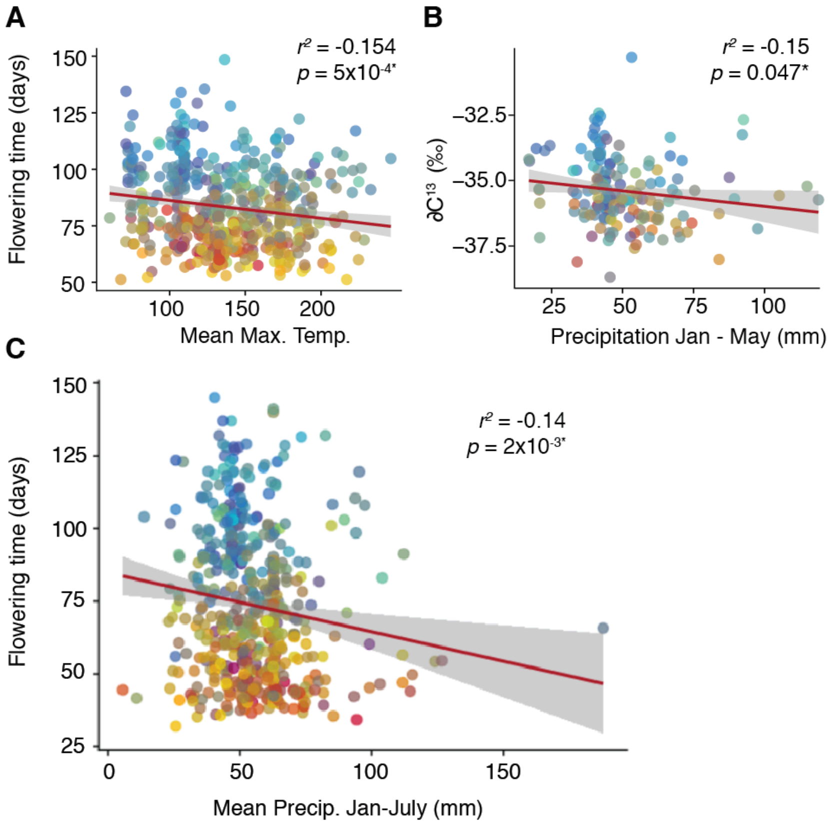


**(A)** Association between flowering time at 16°C and mean maximum monthly temperature, estimates from Pearson's correlation coefficient. (**B**) WUE (using $\delta C^{13}$ as proxy) associated with precipitation averaged for the months January through May. (**C**) Flowering time at 16°C associated with mean precipitation for the months January through July. The months cut-offs do not drive the association, as the associations hold up with the yearly averages as well, however it is more to highlight that extending the growing season to the drier month of July does not change this trend.

**Supplementary Figure 3.** Pairwise Pearson's correlation coefficients for 12 focal traits.

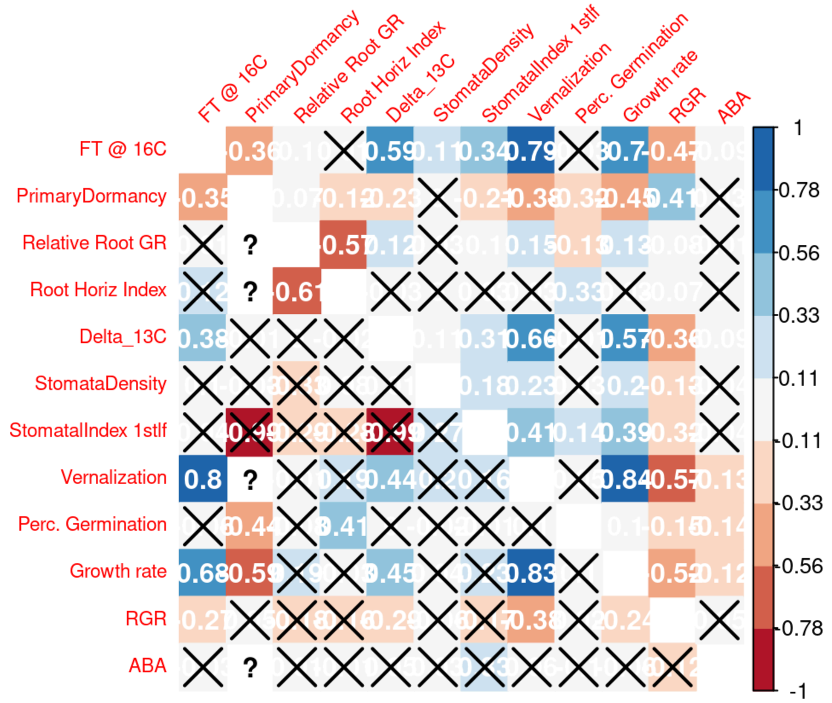


These 12 focal traits were used in the multivariate selection analyses. The upper right triangle are the pairwise correlations amongst imputed trait data and the lower left triangle are the pairwise correlations of the raw trait data. In both, the X indicates the correlation was not significant (p-value > 0.05).

**Supplementary Figure 4.** Total selection coefficients vs. direct selection gradients.

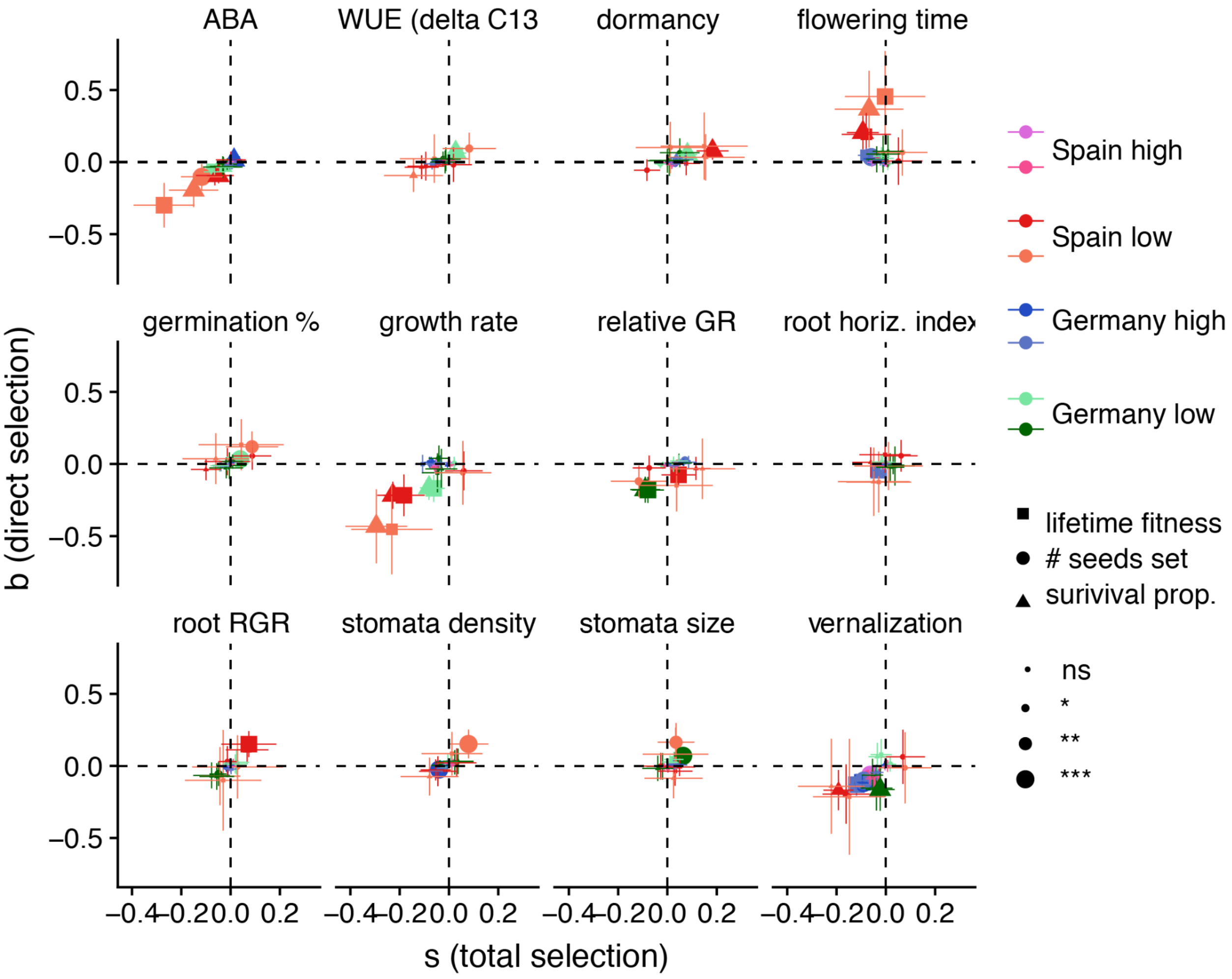


Total selection coefficients (*s*) vs. direct selection gradients ($\beta$, after accounting for trait covariation) and their standard errors for 12 focal traits related to life history and drought adaptation. Estimates made using all three fitness data measurements (lifetime fitness, seeds set, and survival proportion) from two environments hot, Spain and cool, Germany, and in both locations high and low rainfall was simulated. The size of circles indicate significance of $\beta$ and *s* (P < 0.05, 0.01, 0.001).

**Supplementary Figure 5.** Total selection coefficients vs. direct selection gradients from the Germany, high water site.

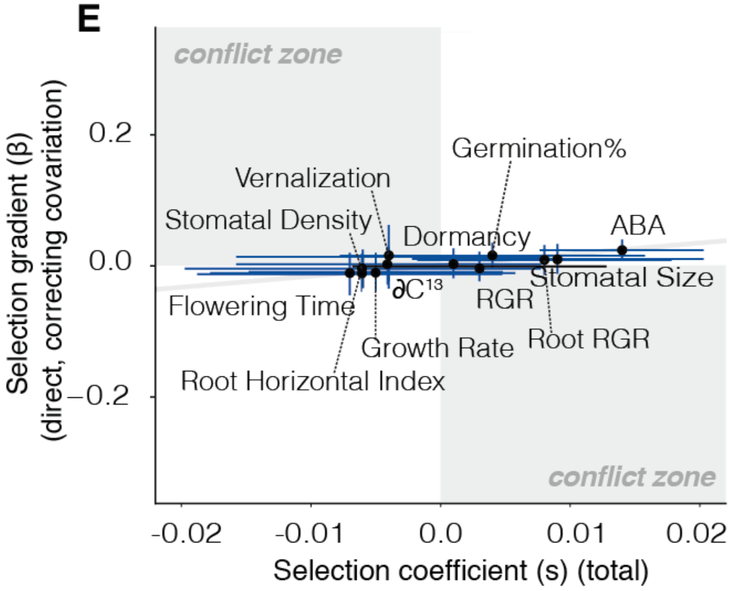


Total selection coefficients (*s*) vs. direct selection gradients (*β*, after accounting for trait covariation), using survival data from the cool and wet common garden experiment (Germany, high water), with high planting density. Large circles indicate significance of *β* and *s* (P < 0.05) and gray areas indicate a conflict where total selection is in one direction (+/-) while direct selection is in the opposite direction.

**Supplementary Figure 6.** Total selection coefficients vs. direct selection gradients without growth rate and vernalization.

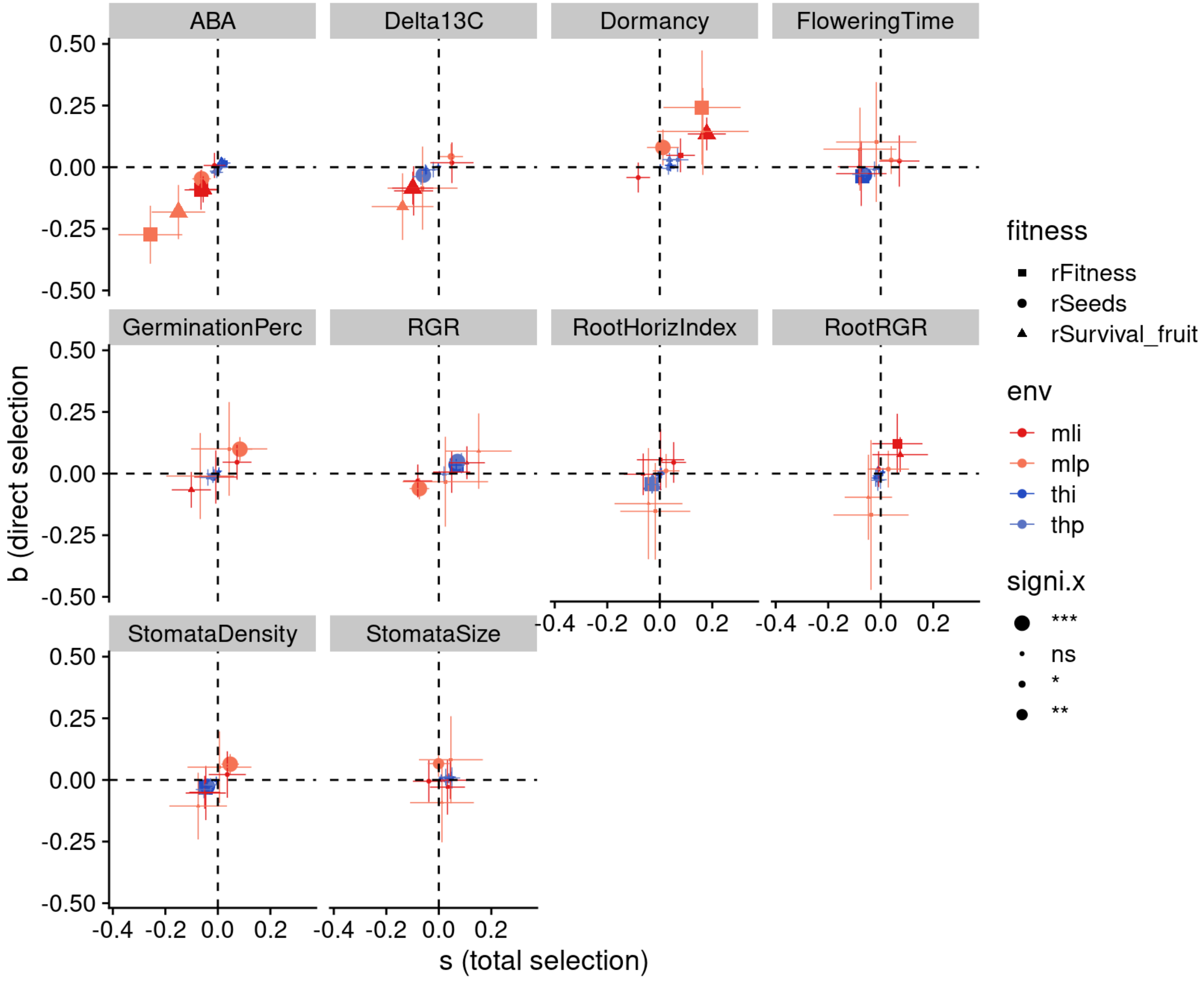


Total selection coefficients ($s$) vs. direct selection gradients ($\beta$, after accounting for trait covariation) and their standard errors for 10 focal traits; notably traits growth rate and growth after vernalization were removed. Estimates made using all three fitness data measurements (lifetime fitness, seeds set, and survival proportion) from two environments hot and dry, Spain (red) and cool and wet, Germany (blue)The size of circles indicate significance of $\beta$ and $s$ ($P < 0.05, 0.01, 0.001$).

**Supplementary Figure 7.** Total selection coefficients vs. direct selection for low rainfall German site.

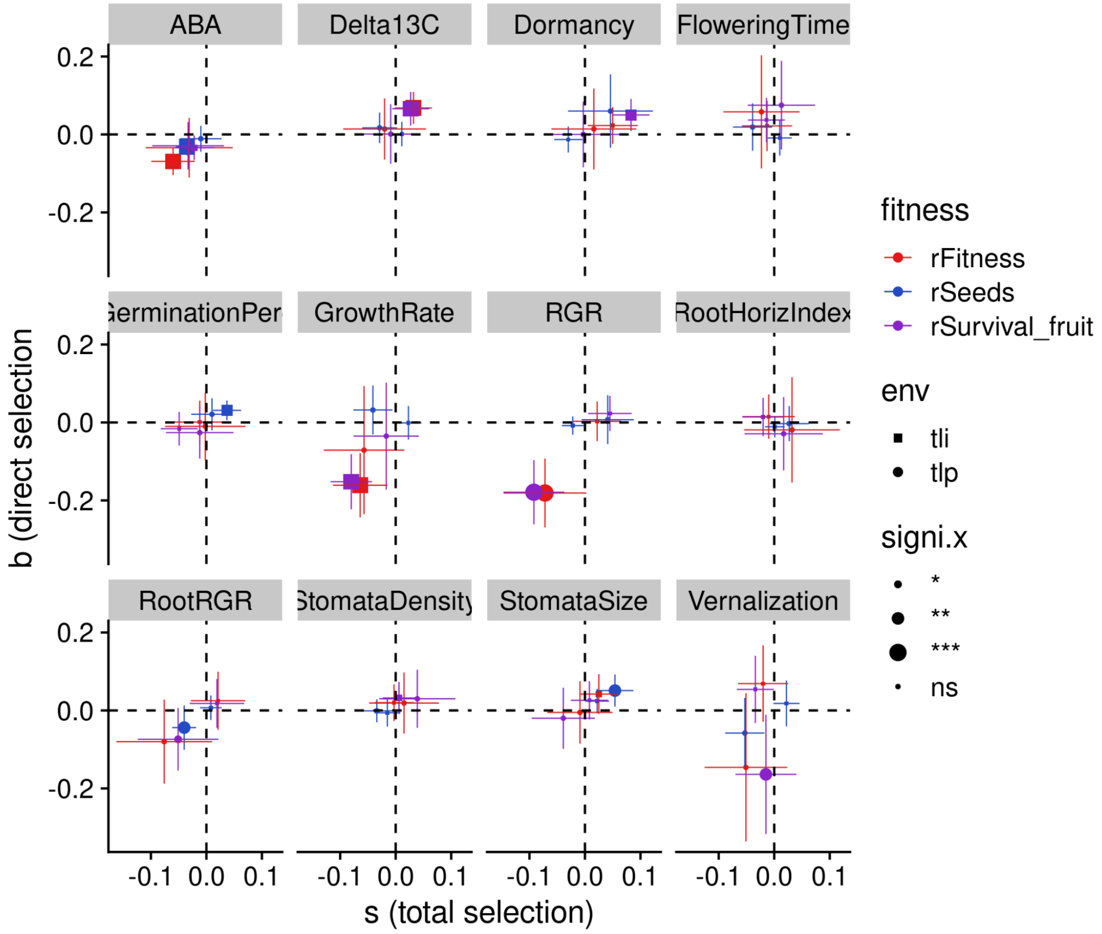


Total selection coefficients ($s$) vs. direct selection gradients ($\beta$, after accounting for trait covariation) and their standard errors for 12 focal traits related to life history and drought adaptation. Estimates made using all three fitness data measurements (lifetime fitness, seeds set, and survival proportion) from the Germany low rainfall site. The size of circles indicate significance of $\beta$ and $s$ ($P < 0.05, 0.01, 0.001$).

**Supplementary Figure 9.** Total selection coefficients vs. direct selection gradients for [26].

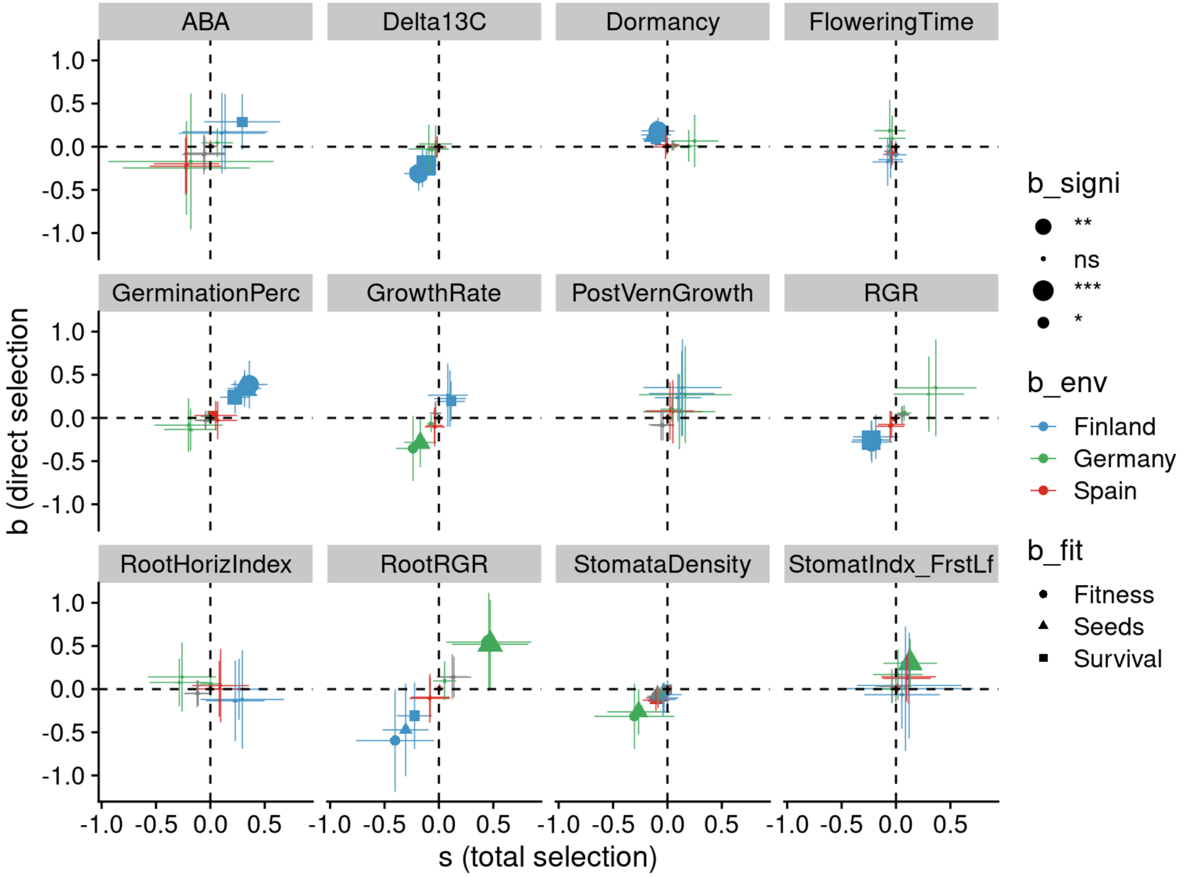


Total selection coefficients ($s$) vs. direct selection gradients ($\beta$, after accounting for trait covariation) and their standard errors for 12 focal traits related to life history and drought adaptation. Estimates made using fitness data (fitness, seeds set, and survival) from three environments in Finland, Germany, and Spain. The size of circles indicate significance of $\beta$ and $s$ ($P < 0.05, 0.01, 0.001$).

**Supplementary Figure 10.** Total selection coefficients vs. direct selection gradients for [27].

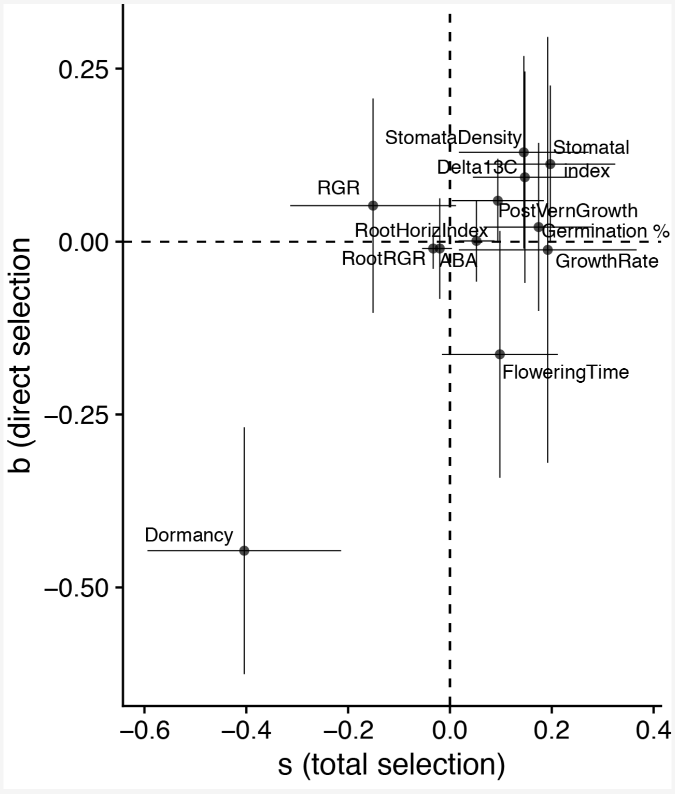


Total selection coefficients (*s*) vs. direct selection gradients (*β*, after accounting for trait covariation) and their standard errors for 12 focal traits related to life history and drought adaptation. Estimates made using fitness and survival data from a common garden site on the Iberian Peninsula [27]. The size of circles indicate significance of *β* and *s* (P < 0.05, 0.01, 0.001).

**Supplementary Figure 11.** Trait correlation data after imputation and normalization.

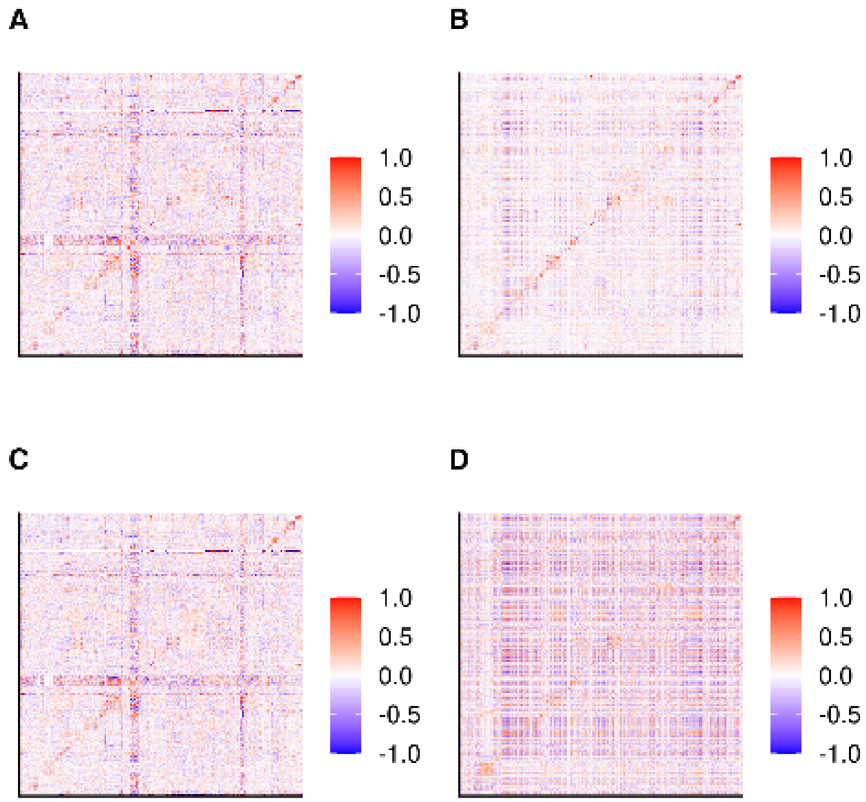


Pairwise Pearson's correlation coefficients for all traits (n=1,862) across various trait datasets A) raw, meaning unimputed and not-normalized B) imputed trait data C) normalized raw data D) imputed normalized trait data.

**Supplementary Figure 12.** Comparison of effect size estimates.

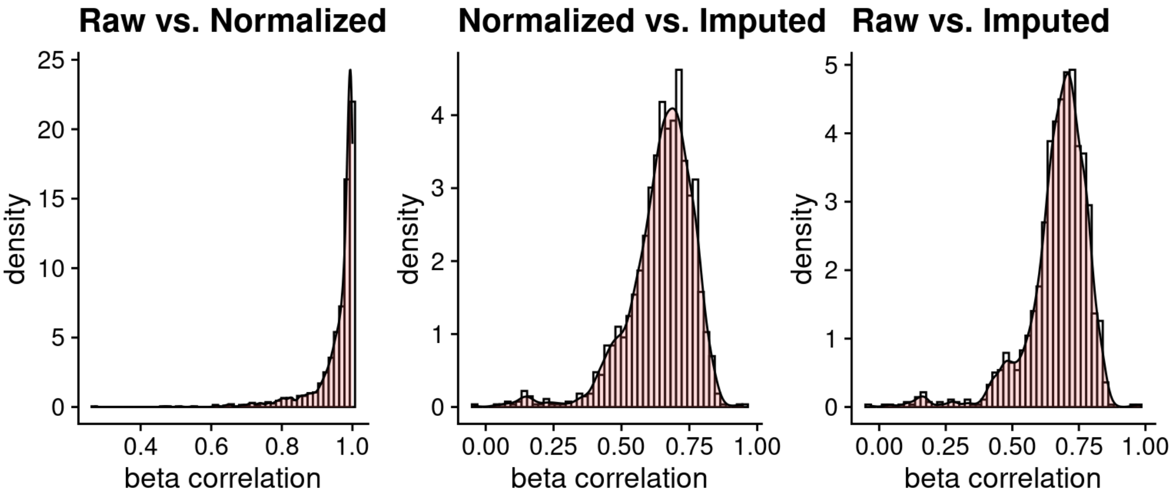


Comparison of 1,862 Pearson's correlation coefficients for effect size estimates from GWA using raw trait data, normalized trait data, and imputed trait data.

**Supplementary Figure 13.** Boxplot of heritability for avoidance and escape traits.

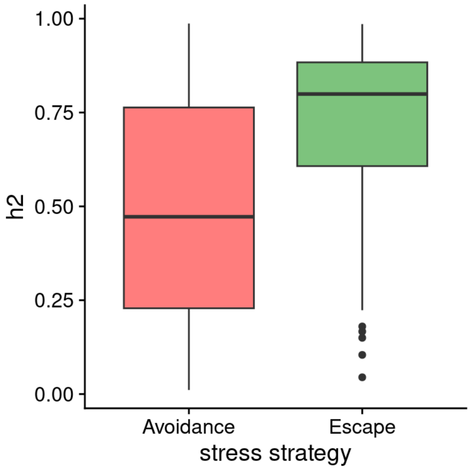


Boxplot of SNP-based heritability estimates for escape traits and avoidance traits post-filtering for traits with inconsistent or low quality heritability estimates; wilcox test p-value=$2.6x10^{-8}$.

**Supplementary Figure 14.** Flowering time and WUE GWA.

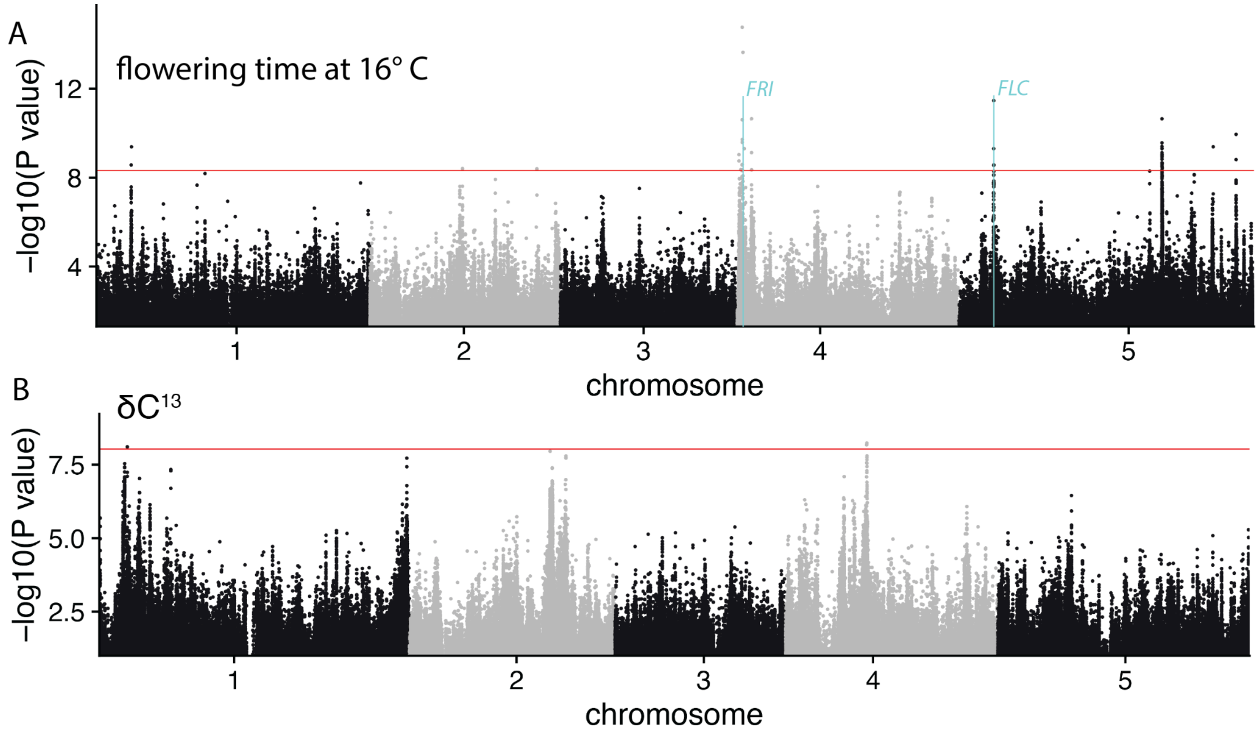


Manhattan plot for (A) flowering time (B) and WUE GWA; data were normalized prior to analysis and not imputed, so only samples with original data were included in each run (n=1021, 261).

**Supplementary Figure 15.** Flowering time and WUE mvLMM GWA.

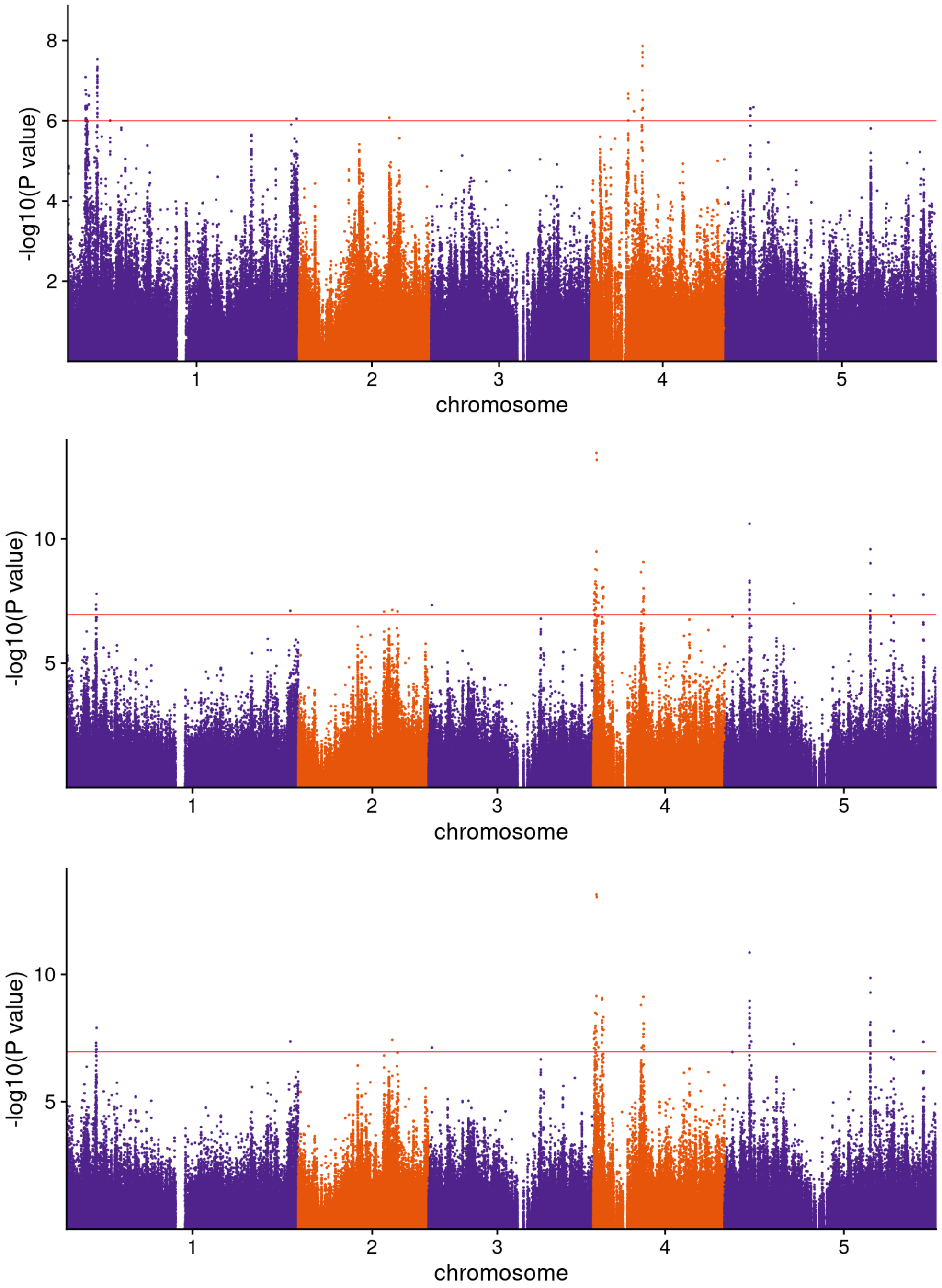


Manhattan plot of mvLMM GWA of flowering time and WUE, corrected with kinship, and with **top**) non-imputed trait data and 5 genetic pcs used as covariates in the analysis, **middle**) imputed trait data, no genetic PCS, and **bottom**) imputed trait data and 5 genetic PCs used as covariates. Note the peak on Chr 1 remains significant and narrows as we correct for population structure with 5 pcs, and this more narrow peak includes only 1 gene on chromosome 1, AT1G11540.

**Supplementary Figure 16.** QQplots for flowering time and WUE mvLMM GWA.

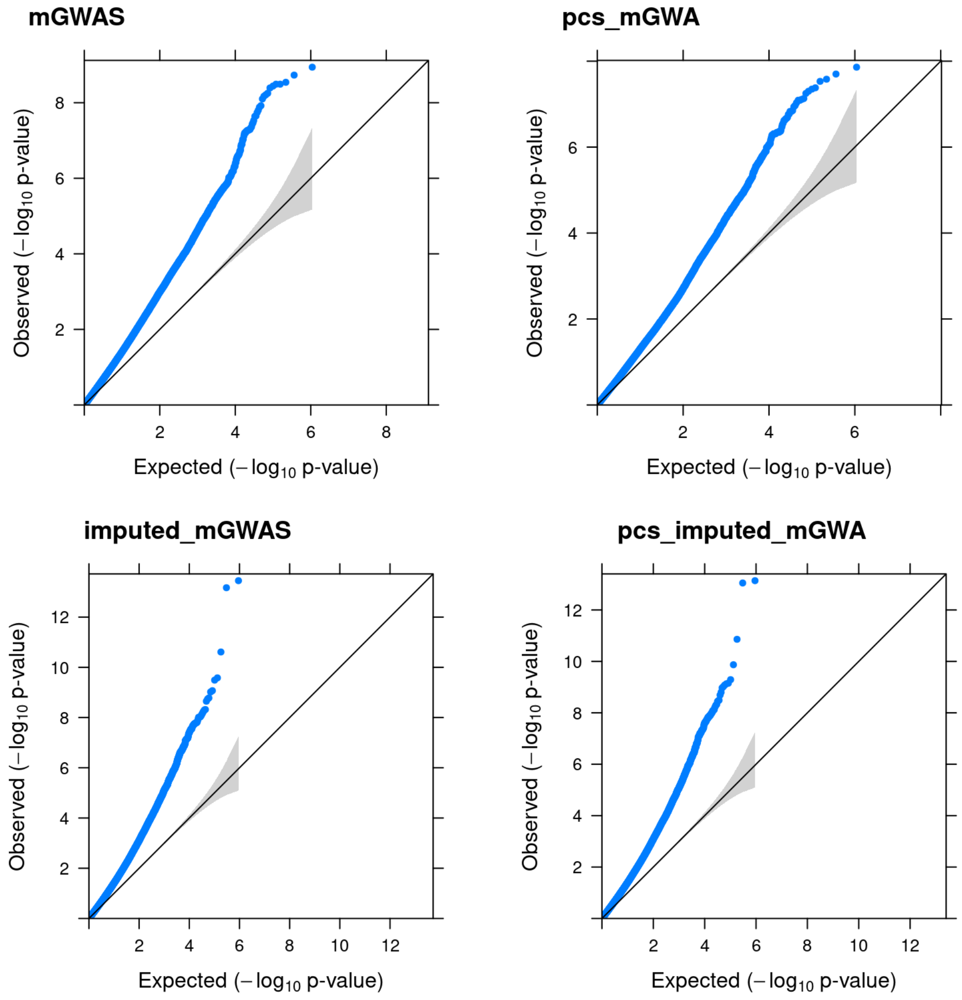


QQplots for mvLMM GWAS with flowering time at 16°C and WUE (δ$C^{13}$) for different variations of both the data and covariates used. (top left) Non-imputed data, n = 248 overlapping accessions. (top right) Non-imputed data, n = 248 overlapping accessions with 5 genetic PCs as additional fixed covariates in the model. (bottom left) Imputed data, n= 1,135 overlapping accessions. (bottom right) Imputed data, n= 1,135 overlapping accessions with 5 genetic PCs as additional fixed covariates in the model.

**Supplementary Figure 17.** Flowering time and growth rate mvLMM GWA.

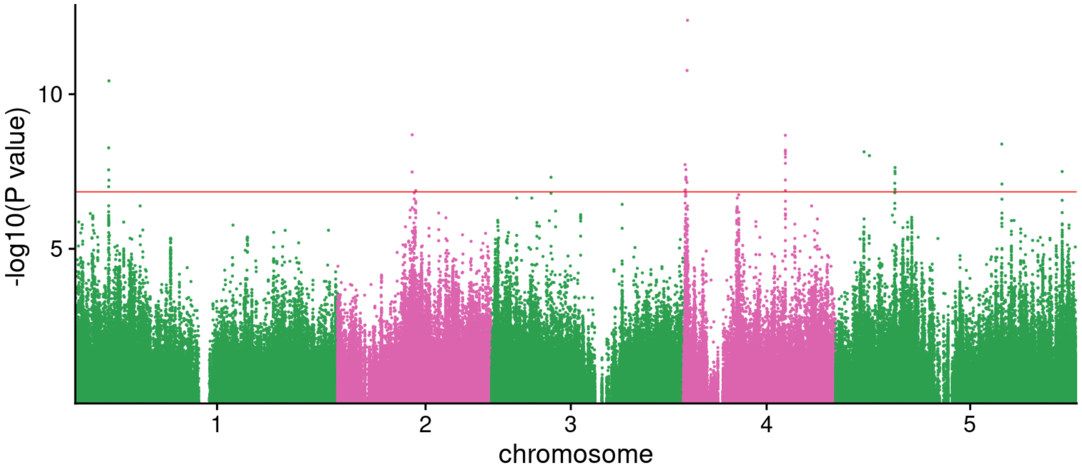


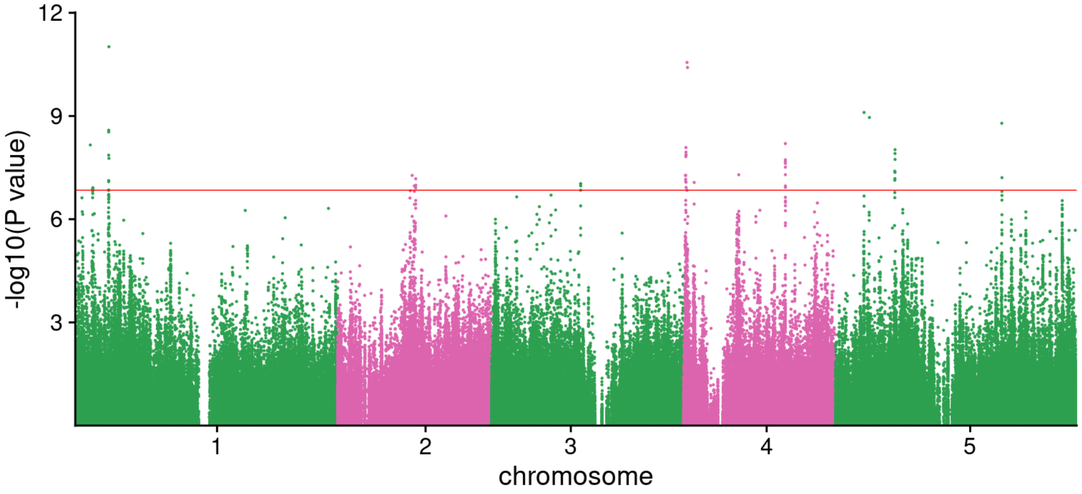


mvLMM GWA between flowering time and growth rate; normalized phenotypic data but not imputed, so the number of overlapping samples, or the total number used in this GWA was 395. Kinship was used to correct for population structure, no genetic PCs (upper), with 5 genetic PCs (lower).

**Supplementary Figure 18.** Genetic correlation estimates between key traits.

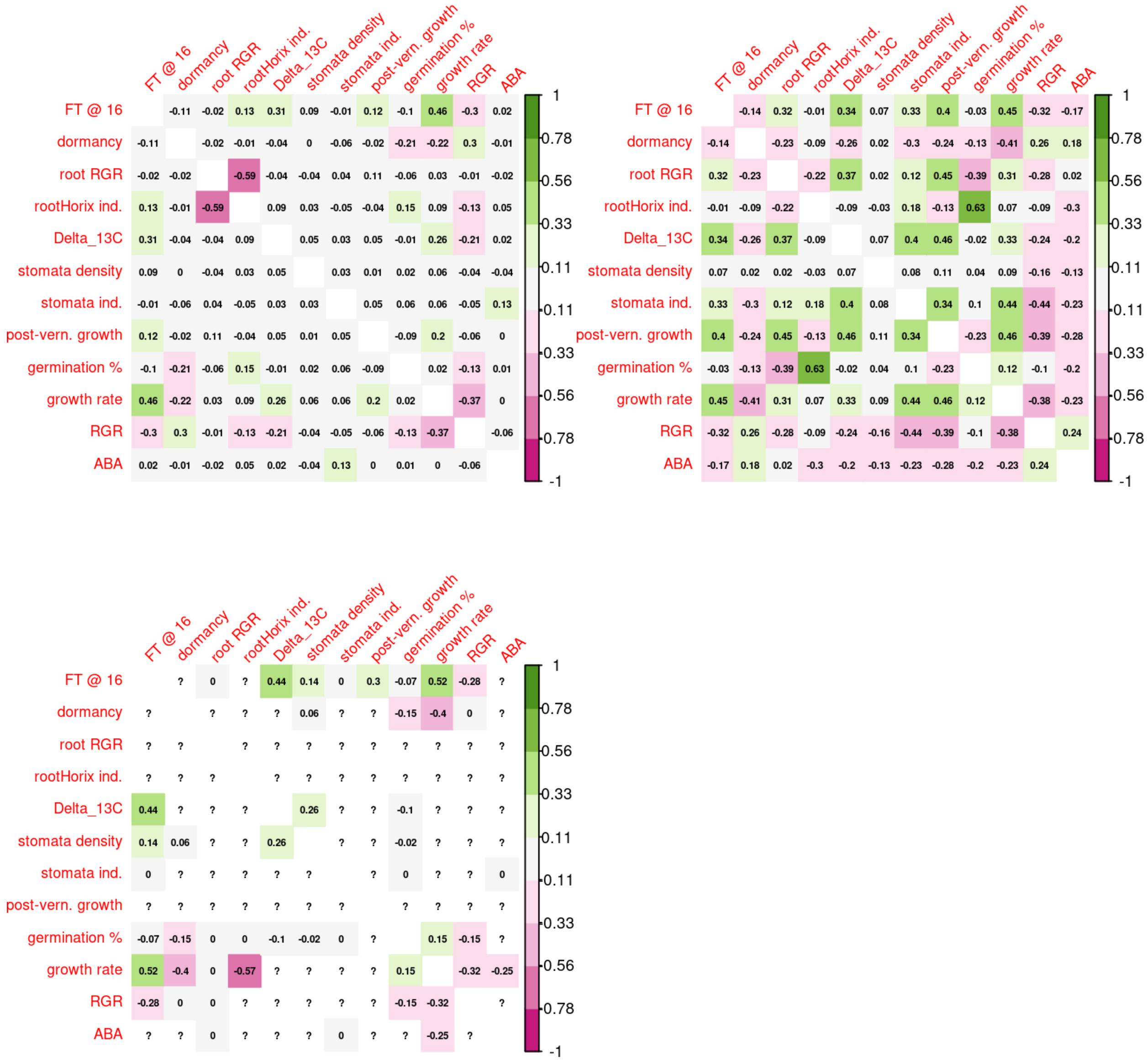


Genetic correlations for 12 key traits estimated through **(top left)** summary statistics for SNPs from independent LD blocks, also called an approximate genetic correlation, **(top right)** imputed and normalized traits used in mvLMM GWA, and **(bottom left)** normalized traits, not imputed, used in mvLMM GWA, in this case many traits did not have enough overlapping samples to perform the method, “?” and “0” indicates not enough data to run the algorithm. All matrices have mirrored top and bottom triangles.

**Supplementary Figure 19.** Flowering time and WUE chromosome 1 peak.

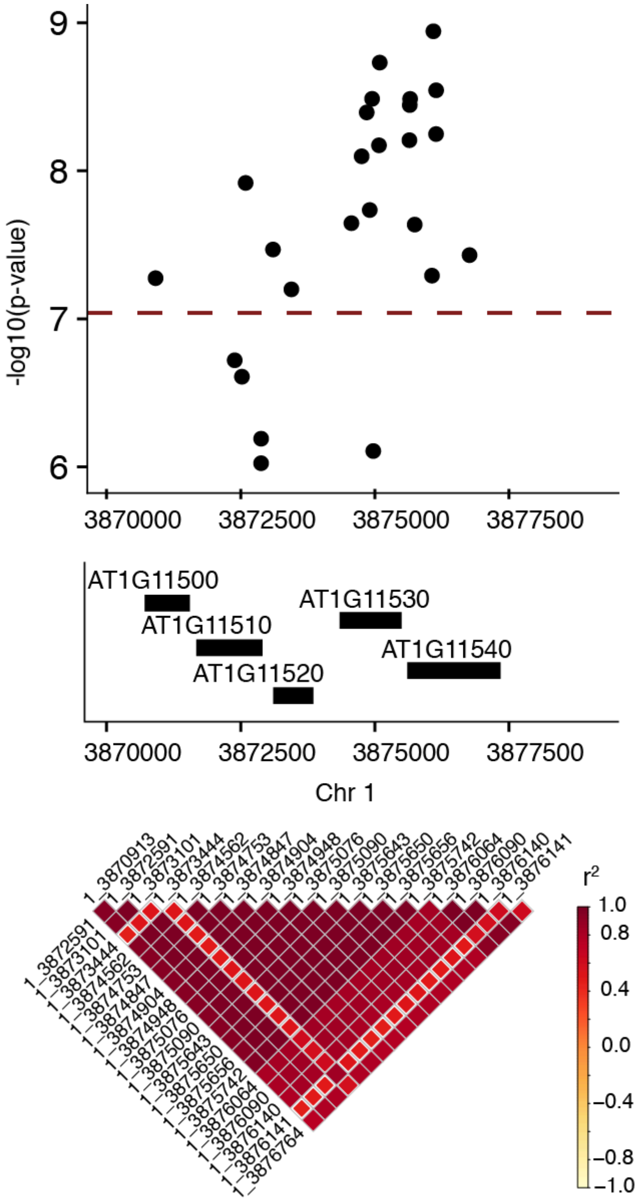


Zoomed in plot of mvLMM GWA for flowering time and WUE effect size peak on chromosome 1 showing the high effect SNPs, the 5 genes in the 7.5kb region and the LD (linkage disequilibrium as esteemed by $r^2$) between the chromosome 1 top SNPs.

**Supplementary Figure 20.** Association of gene expression with key traits.

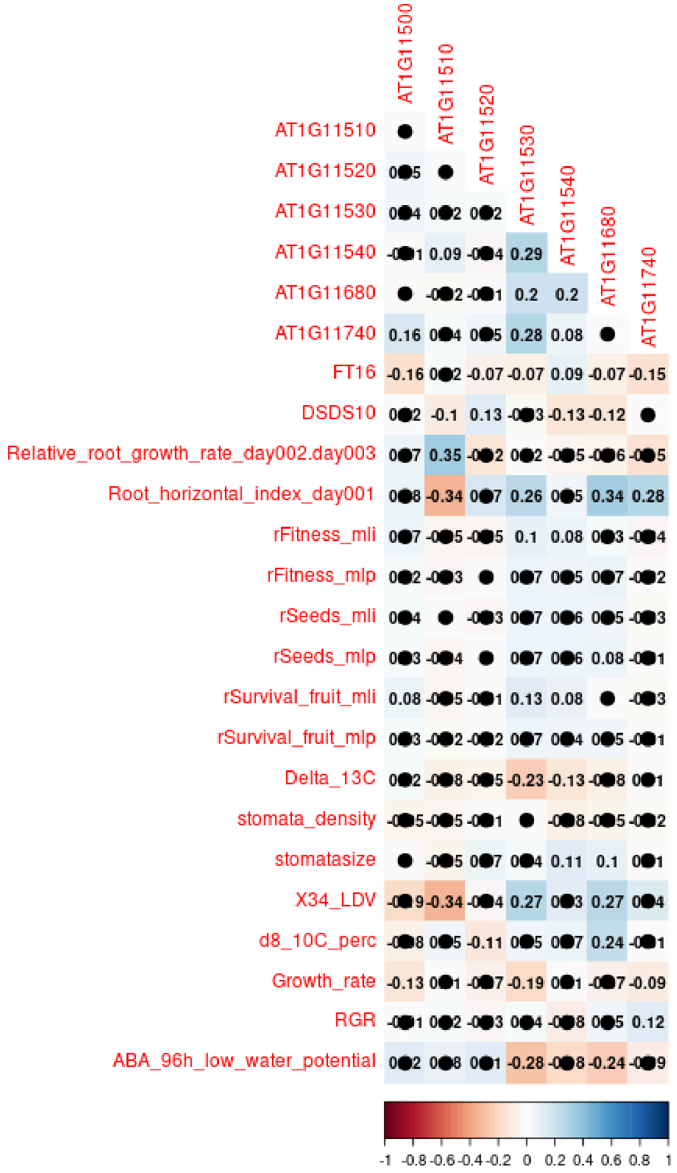


Spearman rank correlations between 12 key traits, a few select fitness data measures, and the expression of the 5 genes identified in the high effect peak on chromosome 1. Cells without a black dot indicated the p-value < 0.2, but note only consider the correlation significant at p-value < 0.05.

**Supplementary Figure 21.** Prediction of trait response to selection.

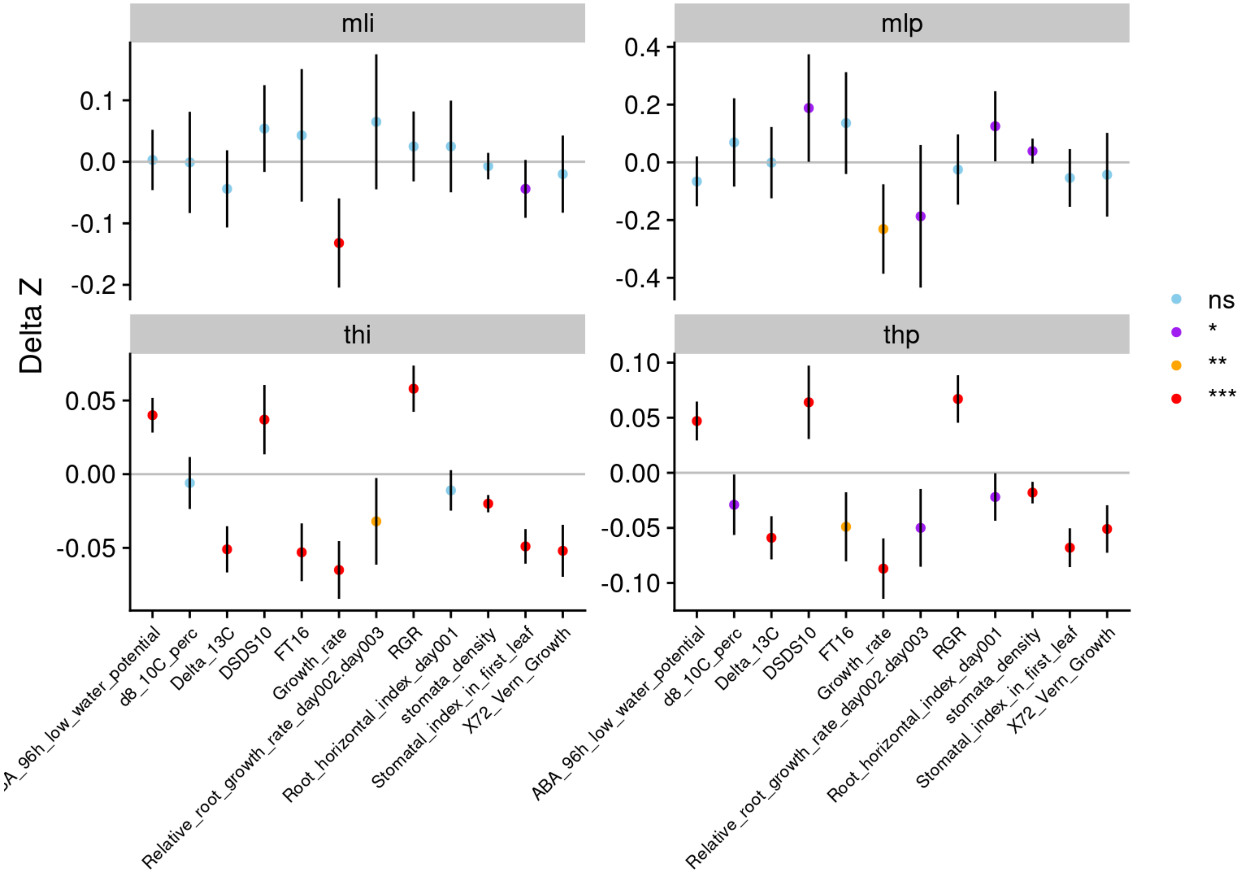


Estimated trait multivariate response to selection in two environments Spain (m) and Germany (t), using trait correlations, genetic correlations estimated from mvLMM GWA, SNP-based heritability, and total selection coefficients. Estimates were made from 100 bootstrap samples, establishing the confidence intervals and significance levels; ns= not significant,*=0.05, **=0.01, ***=0.001.

**Supplementary Figure 21.** Gene editing of diverse *Arabidopsis* accessions.

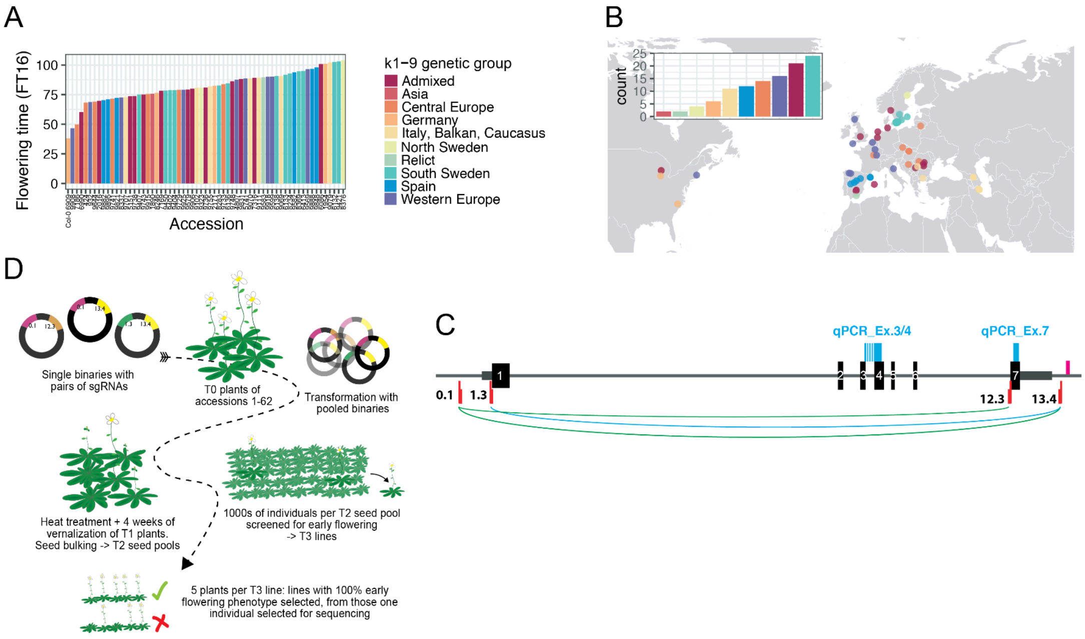


(A) Published flowering time data as days until the first open flower of 62 accessions and Col-0 [1], dataset FT16 (16°C and LD photoperiod [16 h light / 8 h dark]). (B) Geographic origin of the selected accessions. The inset graph shows the number of accessions per genetic group [1]. (C) Schematic representation of the CRISPR/Cas targeted forward genetics approach. (D) Schematic representation of the *FLC* locus. Black boxes represent exons, gray lines show introns, and dark grey indicates UTRs. The blue markings at the bottom illustrate the CRISPR target sites, with the thinner part representing the position of the respective PAM site. Two of the selected Cas9 target sites were located upstream of the FLC translational start site: one in the promoter region (site 0.1, -349 bp relative to the A of ATG) and one in the 5' UTR region (site 1.3, -19 bp). Downstream target sites are located in exon 7 (site 12.3, -5518 bp) and the 3' downstream region (site 13.4, -6055 bp). The curved lines indicate the pairwise combinations of target sites, as they were cloned as single binary vectors. The blue bars indicate the binding sites of the two pairs of qRT-PCR primers.

**Supplementary Figure 22**. *FLC* sequence coverage across accessions.

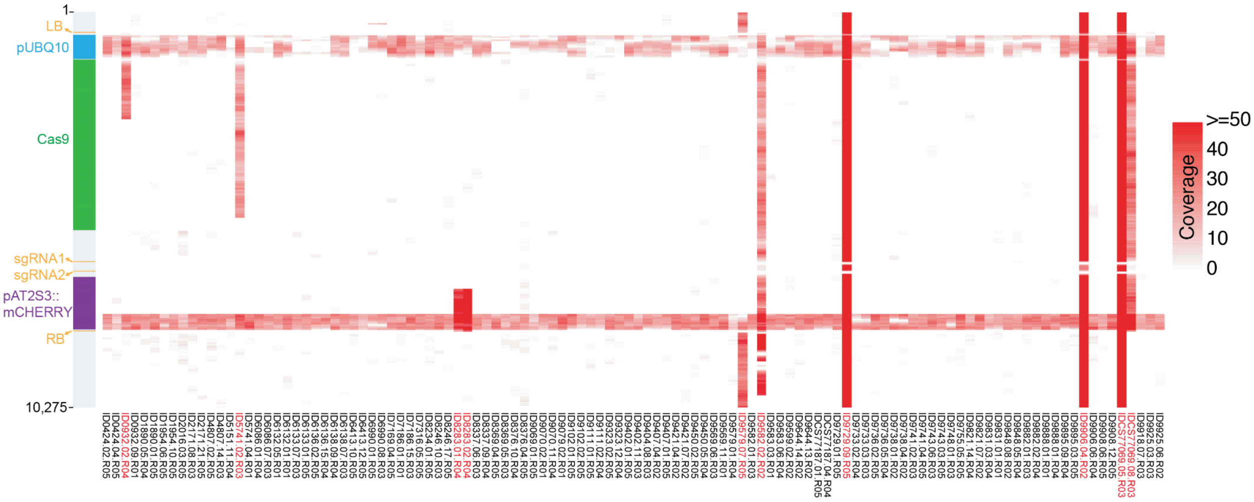


Reads of whole-genome sequencing (WGS) with a mean genome-wide coverage of 26.5x (min: 17.2x, max: 43.0x) from each line were mapped to the supermodule expression plasmid [41]. One column per line. The color code indicates the coverage 0 to 50 or higher. The bar on the left illustrates the different supermodule elements. Ten of the 112 lines (9%) had one to four insertions of portions of the transgene that were not derived from *A. thaliana* sequences (marked with red line IDs, **Table S25**).

**Supplementary Figure 23.** Relative expression of *FLC* across accessions.

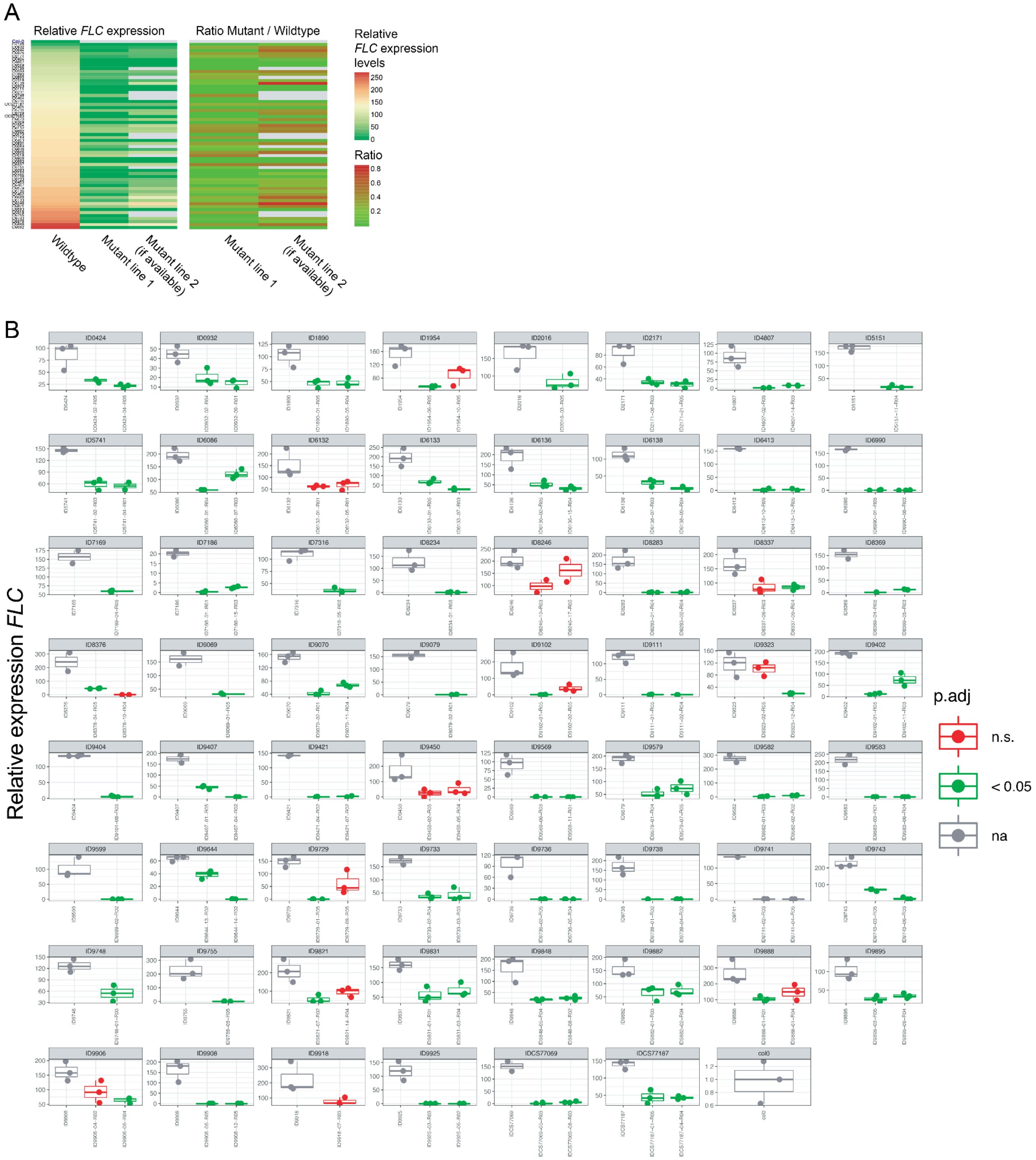


(A) Transcript analysis of *FLC* via qRT-PCR from nine-day-old plants grown under long days. On the left, the relative *FLC* expression data is shown. All samples were sorted by the expression levels of the wild types, and one or two of the corresponding mutants are shown. Most of the 112 lines, 94 (84%), representing 57 accessions, had reduced amounts of *FLC* transcript, from 12.9% to 100% (Student's t test, Benjamin-Hochberg correction, p.adj < 0.05). 39 lines representing 26 accessions are likely complete knock-outs, and the remainder are knock-down (KD) lines. KD and KO alleles in the same genetic background were obtained for nine accessions (**Table S27**). On the right, the ratio of the relative expression levels between each wild type and its corresponding mutant lines is shown. (B) The data used to generate A. The color code indicates the p-value of a two sided Student's t test with Benjamini-Hochberg correction of each mutant versus the respective wild type. The relative quantification was calculated with the ΔΔCt method using ACT8 (AT1G49240) as a standard [55] and calibrated by biological replicate 1 (rep1) of Col-0.

**Supplementary Figure 24.** Comparison of relative expression across exons.

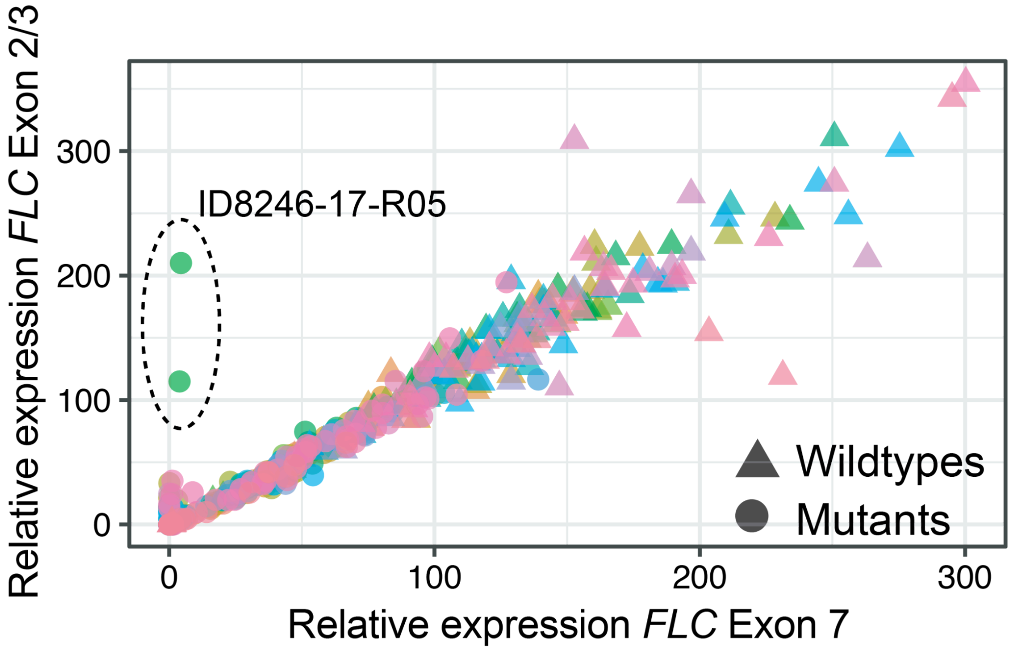


Comparison of the *FLC* transcript levels as measured by primers targeting exon 2/3 or exon 7. The line ID8246-17-R05 (two biological replicates, circled and labeled in the graph) of accession 8246 expressed a truncated transcript, which is most likely non-functional.

**Supplementary Figure 25.** Summary of δ13C data from mutants and wildtypes.

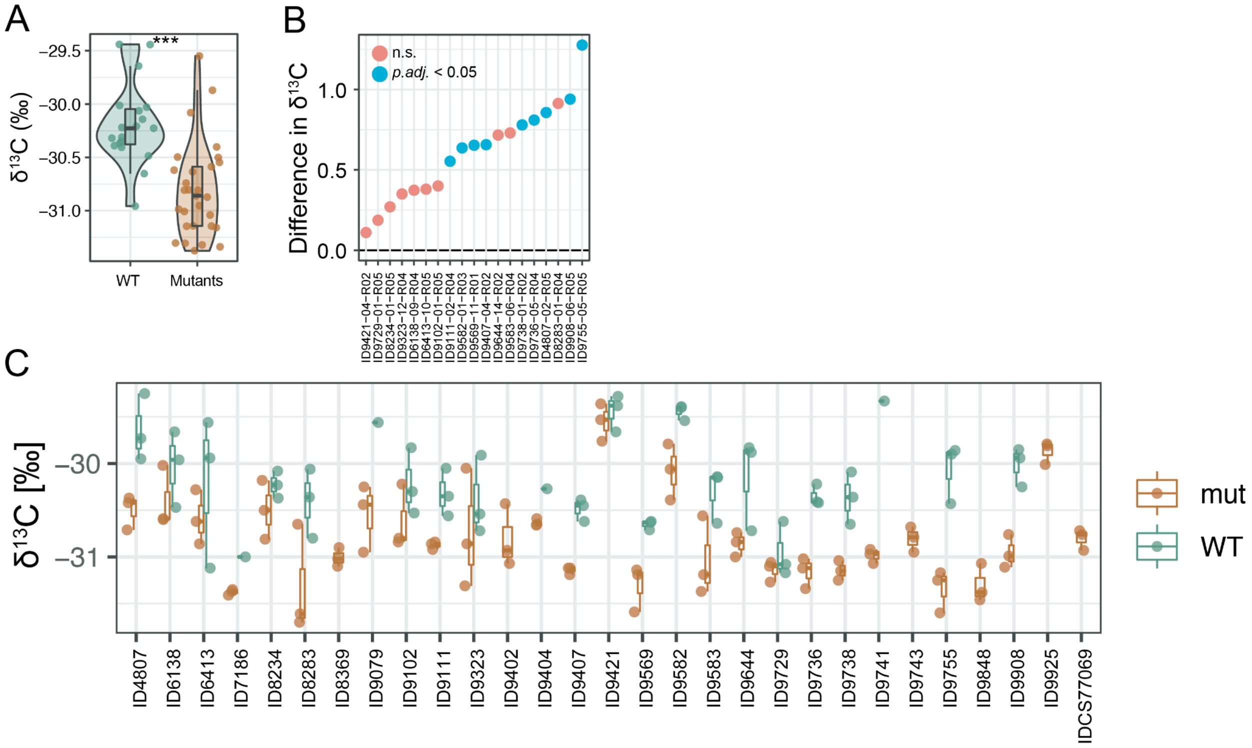


(A) Carbon isotope composition (δ13C [‰]) 20 wild-type accessions (including Col-0) and 29 flc mutants (with a maximum relative FLC expression level of 18.3 a.u.), which included 19 flc mutant / wild type contrasts. The mean values of three biological replicates are shown. Wild-types, N=19; mutants, N=29. Mean [±se] mutants: -30.81‰ [±0.082]; mean wildtypes: -30.19‰ [±0.087]; Mann-Whitney U rank test, p=7.0e-06. (B) Difference in $\delta^{13}C$ ($\delta^{13}C^{diff} = \delta^{13}C_{Wildtype} - \delta^{13}C_{Mutant}$) between wildtype and the respective mutant. Blue dots indicate a significant difference, two-sided Student's t-test, Benjamini-Hochberg correction, p.adj. < 0.05. (C) δ13C [‰] of three biological replicates per line. Results of the pairwise comparisons with the respective wild type shown in C (two sided Student's t test, Benjamin-Hochberg correction, p.adj. < 0.05).

**Supplementary Figure 26.** Correlation of mutant *flc* δ13C [‰] values and those from [19].

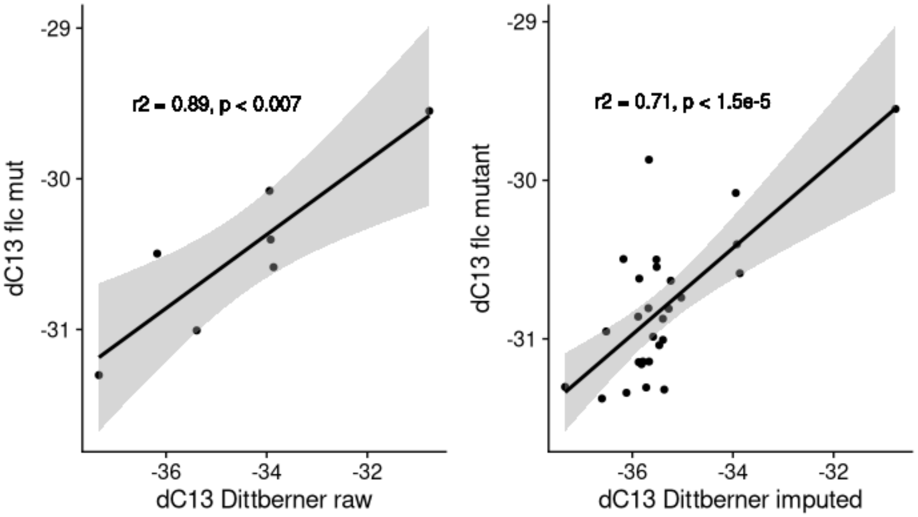


Left) Correlation between *flc* mutant δ13C [‰] and accession used in [19], note there are only 6 true overlapping accessions. Right) correlation between the imputed δ13C [‰] data for the 1001 Genomes Accessions that was imputed in this study (see **Text SI**).
